# Multimodal conformational landscape and thermodynamics of coupled folding and binding in an intrinsically disordered protein region

**DOI:** 10.64898/2026.11.13.766518

**Authors:** Vasily Akulov, Valerio Rizzi, John van Noort, Alireza Mashaghi

## Abstract

Intrinsically disordered proteins and peptides, as well as flexible protein regions, often bind folded protein partners through coupled folding and binding, but the underlying conformational landscapes and thermodynamics remain challenging to resolve. The large conformational space of the unbound molecule and its interactions with the folded partner make it difficult to identify and efficiently sample the slow collective motions governing association and folding. Here, we use enhanced-sampling molecular dynamics to investigate the coupled binding and folding of the intrinsically disordered region of FCP1 phosphatase upon interaction with the folded transcription factor Rap74, a process that modulates the activity of RNA polymerase II. OneOPES enhanced sampling simulations reveal a multimodal binding landscape comprising native, flipped and disordered-bound states, and identify hydrophobic contacts as early determinants of association followed by progressive helix formation. Circuit topology analysis further elucidates the FCP1 folding mechanism by identifying a characteristic topological transition accompanying helix formation and its stabilisation upon binding. To resolve the thermodynamics of the process, we introduce *Divide et impera*, a thermodynamic decomposition that separates the free-energy cost of folding from the binding free energy of the folded conformation. This approach reduces the conformational complexity of individual calculations while preserving the thermodynamic connection between unbound and bound states, enabling the calculation of absolute binding free energies in close agreement with experiment. These results establish a framework for connecting conformational heterogeneity, binding mechanisms and thermodynamics in coupled folding and binding of disordered proteins, with potential applications in the rational design and development of peptide therapeutics.

## Introduction

Protein binding underlies a wide range of biological processes and is traditionally viewed as an association between two preformed, stably folded structures(Rapti et al., 2026). However, increasing attention has been directed towards interactions involving flexible and dynamic protein regions, such as the variable regions of antibodies and T-cell receptors. A particularly important class of interactions occurs when an intrinsically disordered protein or region (IDP/IDR) binds to a folded partner while undergoing substantial conformational rearrangement. Unlike folded proteins, IDPs do not adopt a single compact structure but instead populate heterogeneous ensembles of rapidly interconverting conformations (Akulov et al., 2025). Upon interaction with a binding partner, some IDPs undergo substantial conformational rearrangement and form stable, structured complexes. The resulting binding process can involve a broad ensemble of transient conformations and competing pathways, making it difficult to determine how molecular interactions drive the transition from the disordered ensemble to the bound state, and to quantify the associated binding free energy. Resolving these pathways and intermediates and the underlying free energy landscape is important for understanding the molecular basis of IDP-mediated recognition, regulation, and peptide drug development (Löhr et al., 2026).

Recent advances in machine-learning-based structure prediction have greatly expanded our ability to model protein structures and biomolecular complexes. AlphaFold3, for example, can predict structures of complexes containing proteins and a broad range of other biomolecular components (Abramson et al., 2024). Generative approaches such as BioEmu further enable efficient sampling of approximate protein conformational ensembles (Lewis et al., 2025). However, these approaches do not directly describe the thermodynamic landscape or the dynamical pathways connecting states in dynamic proteins. Consequently, they cannot by themselves determine the driving forces underlying conformational transitions or distinguish between alternative binding mechanisms such as conformational selection and induced fit. Moreover, generating accurate conformational ensembles for multicomponent systems remains challenging because the available training data for protein–protein interactions are limited. Molecular dynamics simulations provide an atomistic framework for such analyses, but coupled folding and binding can occur on timescales that are difficult to access with conventional simulations.

Long-timescale atomistic simulations have demonstrated that such sampling is possible but computationally demanding (Chong & Ham, 2019; Robustelli et al., 2020). For example, Robustelli et al. used hundreds of microseconds of atomistic simulation to observe numerous binding and unbinding events for the interaction between the intrinsically disordered C-terminal domain of the measles virus nucleoprotein and the X domain of the phosphoprotein, enabling analysis of the free-energy landscape and binding kinetics. Such extensive simulations remain computationally expensive, motivating enhanced-sampling approaches that accelerate rare conformational transitions while retaining an atomistic description of the system. These methods include metadynamics (Laio & Gervasio, 2008), bias exchange metadynamics (Domene et al., 2015) and infrequent metadynamics (Tiwary & Parrinello, 2013). More recently, the On-the-fly Probability Enhanced Sampling (OPES) method and its OneOPES framework have enabled efficient exploration of complex free-energy landscapes and have been applied to absolute binding free-energy calculations(Invernizzi et al., 2020; Invernizzi & Parrinello, 2020, 2022; Rizzi et al., 2023). Enhanced-sampling approaches have also been used to investigate IDP folding and binding, including p53 peptide binding to MDM2 (Zou et al., 2020) and the simulation of α-MoRE-MeV-XD complex (Han et al., 2016). Together, these developments provide an opportunity to investigate coupled folding and binding without relying exclusively on exceptionally long unbiased trajectories.

Despite substantial progress in characterizing IDP binding, a quantitatively converged free-energy landscape that simultaneously captures binding thermodynamics, structural heterogeneity, and multiple binding pathways has remained elusive. Here, we combine enhanced-sampling simulations, Circuit Topology, and free-energy calculations to characterize the coupled folding and binding of the disordered domain of FCP1 phosphatase (FCP1_term_) to RAP74, the well folded subunit of transcription factor Transcription factor IIF (TFIIF) (Archambault et al., 1998; Nguyen, Chen, et al., 2003). The interaction regulates dephosphorylation of RNA polymerase II, during the transcription cycles, and has been extensively studied both experimentally and computationally (Gibbs & Showalter, 2016; Kamada et al., 2003; Kumar et al., 2013; Lawrence et al., 2011; Lawrence & Showalter, 2012; Nguyen, Abbott, et al., 2003; Wostenberg et al., 2011) leading to key insights into their structure and function, yet a converged atomistic description of the free-energy landscape connecting the disordered, partially folded, and bound states remains lacking. In particular, it remains unclear how individual contacts and secondary-structure formation are coupled during binding and to what extent alternative binding pathways contribute to complex formation.

We first characterize the conformational landscape of FCP1_term_ and its folding transitions in solution and then resolve the structural pathways connecting the disordered and bound states of the complex. Motivated by the thermodynamic treatment of folding as a penalty associated with binding (Hadži & Lah, 2022), we formulate a thermodynamic cycle that separates the free energy of folding from the free energy of binding. We compare this decomposed approach with a direct calculation of coupled folding and binding and assess its convergence and reproducibility. Finally, we examine alternative binding pathways to distinguish features consistent with induced-fit from conformational-selection mechanisms. Together, these analyses provide an atomistic framework for connecting conformational ensembles, binding mechanisms, and thermodynamics in coupled folding-and-binding processes.

## Methods

### Molecular Dynamics Simulations

The interactions between Rap74 and FCP1_term_ were investigated with a combination of molecular dynamics and circuit topology. First, to study the conformational ensemble of FCP1_term_ in isolation, we simulated it using conventional and OneOPES (Rizzi et al., 2023) enhanced sampling protocols. Conventional MD simulations were performed for 10 µs at 300K. We performed four independent OneOPES simulation repeats, each comprising eight replicates; each replica was simulated for 500 ns. We then performed contact and circuit topology analyses, and compared the obtained ensembles with previously published NMR data.

Next, we analyzed stability of the Rap74:FCP1_term_ complex using conventional equilibrium MD simulations at 300 and 350 K, each with a duration of 10 µs. We then studied the mechanism of complex formation. To investigate the conformational space of the complex, we performed exploratory enhanced sampling simulations using OPES_EXPLORE method. The bias was applied along two collective variables, LDA_interchain_ and LDA_helix_, the first tracked native interchain contacts and the second one the native helical contacts within the FCP1_term_. We performed four independent simulation repeats, each lasting 8 µs.

The resulting ensembles were used to adapt biasing CVs. The new main CVs were tracking native and non-native interchain contacts, and the distance between FCP1_term_ center of mass and binding pocket on the surface of Rap74. With these CVs, we studied complex formation. We did it using two types of OneOPES simulations: in the first the folding of FCP1t_erm_ were restricted to resemble the native helical formations, in the second no such restriction was applied. Instead, the helical content of FCP1_term_ was included as an additional CV. This allowed us to compare different folding and binding mechanisms including induced fit and conformational selection, and calculate the binding free energy. We performed four independent repeats of each OneOPES protocol, each comprising eight replicas. Each replica was simulated for 500 ns.

All simulations were performed using GROMACS 2023.5 patched with the PLUMED 2.9.3. A detailed description of the methods is provided in the supporting materials (Abraham et al., 2015; Tribello et al., 2014).

### NMR

The NMR chemical shifts were calculated from the simulated trajectories using the SPARTA+ program (Shen & Bax, 2010). The experimental NMR spectrum was obtained from (Showalter & Showalter, 2009). We corrected the NMR backbone chemical shifts by subtracting the corresponding random coil component from both the predicted and calculated spectra. This component was calculated using the Poulsen IDP/IUP random coil chemical shift server (Kjaergaard & Poulsen, 2011, 2012).

### Circuit topology

We used circuit topology (CT) analysis to characterize the organization of intrachain contacts during conformational changes in FCP1_term_. CT considers pairs of intrachain contacts and classifies their arrangement into three topological types: series (S), in which the two contacts occur sequentially along the chain; parallel (P), in which one contact is nested within the other; and cross (X), in which the two contacts cross along the chain(Golovnev & Mashaghi, 2020; Hammond et al., 2026; Heidari et al., 2020; Mashaghi & Ramezanpour, 2015b, 2015a). For each conformation, we quantified the corresponding topology using the ProteinCT software (Moes et al., 2022). The resulting topological arrangements were visualized using PyMOL with the circuit topology plugin (Dimiņš et al., 2025).

## Results

### FCP1_term_ is predominantly disordered but transiently folds into an α-helical conformation

First, we performed a 10 µs conventional MD simulation of FCP1_term_ in an aqueous solution. We observed that FCP1_term_ transiently forms an extended α-helical structure (Figure 1A) as well as some short-lived β-sheet structures. The helical structure is particularly interesting, as this conformation forms when FCP1_term_ interacts with Rap74. Consistent with previous results, we find that the most stable state of FCP1_term_ is disordered: 82.4% of frames have a value of the helical-turn collective variable, S_helix_, below 3. As can be seen in Figure 1A, FCP1_term_ has fewer than 3 residues in a helical conformation for most of the time. However, at 5.8 µs and 9.8 µs, more than eight residues adopt a helical conformation, corresponding to 6.4% of the trajectory frames. When this extended helical structure forms, fluctuations in the radius of gyration decrease.

**Figure 1.**
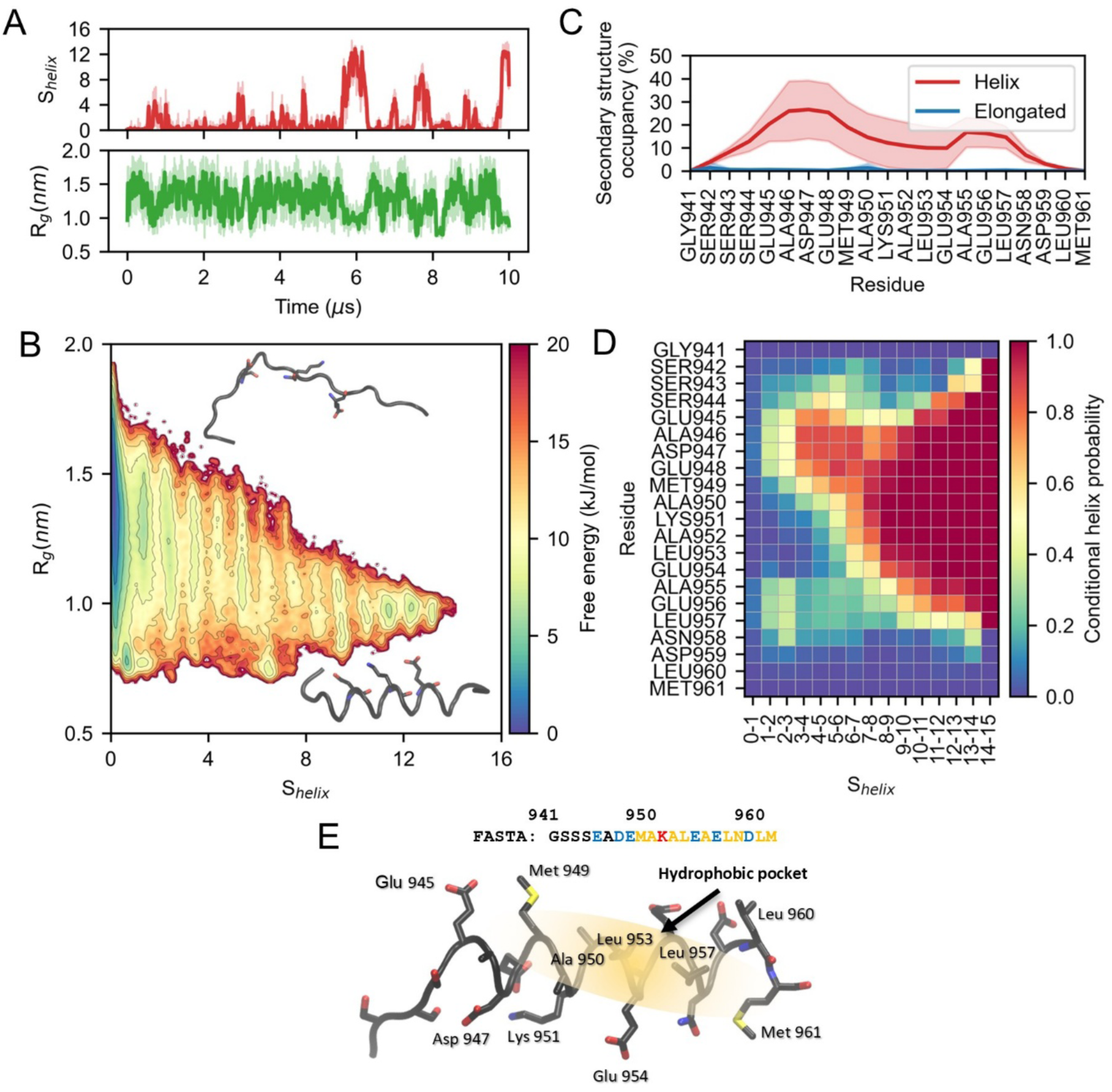
FCP1term is predominantly disordered but exhibits a folding propensity driven by charged residues. (**A**) The dynamics of the helical turns collective variable (top) and radius of gyration (bottom) during the unbiased simulation of the isolated FCP1term. (**B**) The free energy surface of FCP1term (fragment of RNAP II CTD Phosphatase) as a function of the radius of gyration Rg and helical turns collective variables. (**C**) Secondary structure occupancy for FCP1term, determined from conventional simulation, calculated with DSSP (Hekkelman et al., 2025). The red line represents helical structure, the blue line represents elongated structure. Shading represents the standard deviation, calculated from the block analysis, 5 blocks were used. (**D**) 2D conditional helix probability map. The abscissa shows ranges of the helical collective variable bins, Shelix, and the ordinate shows residue numbers and amino acid identities. The color represents the probability that a given residue adopts a helical conformation when the overall helicity, Shelix, falls within the corresponding range. (**E**) The sequence and folded structure of the FCP1. The yellow area shows the hydrophobic patch.

In Figure 1B, using the same helical collective variable and radius of gyration as reaction coordinates, we constructed a 2D free-energy surface (FES). The high-helicity state is approximately 8 kJ/mol higher in free energy than the most stable disordered state. Thus, the extended helical conformation is relatively unfavourable but remains accessible during the simulation.

Different residues exhibited different propensities to form an α-helical structure (Figure 1C). The N- and C-terminal residues are predominantly disordered, whereas two regions exhibit higher helical propensity, i.e., between residues 942–951 and 954–960. The first helical region has a helical propensity of approximately 25%, while the second is around 10%. However, the block analysis shows much larger fluctuations in the first region, which could reflect slow convergence of the simulation statistics. Therefore, the actual helical propensity of this region may be closer to that observed in the independent simulations described below.

To understand the mechanism of helix formation, we constructed a 2D histogram of the conditional helix probability (Figure 1D). For each number of helical turns, we calculated which residues were most frequently present in the helical conformation. This allowed us to estimate the probability of each residue adopting a helical conformation as a function of the total helicity of FCP1_term_. The resulting plot shows a strong contrast between the residues involved in short and long helical structures. When the total number of helical turns is below 7, in agreement with Figure 1C, the helical structure is mainly formed around residues 942–951 and 954–960. However, for the formation of a long and stable helix (*S*_helix_ > 7), incorporation of Lys951 into the helical structure appears to be necessary.

We therefore suggest that the formation of the extended helix may be stabilised by electrostatic interactions involving Lys951 and the surrounding negatively charged residues Glu947 and Glu954 (Figure 1E). At the same time, formation of the extended helix exposes a hydrophobic surface containing Leu residues, which favours helix formation and leads to a decrease in interactions with the solvent. However, Met949 approaching this hydrophobic cluster creates tension that can break the helical structure. Another destabilisation factor is the change at the N-terminus of the peptide. It makes contacts with Glu471 and repels Lys951.

We validated the conformational mechanism observed in conventional MD simulations using four independent OneOPES simulation repeats (Figure S1). The primary CV used in these simulations was S_helix_, which quantifies the number of residues adopting a helical conformation in the FCP1_term_ peptide. All four repeats sampled a fully folded helical structure within 200 ns. The conformational landscapes of the four repeats showed a characteristic triangular distribution in the R_g_–S_helix_ space. This distribution was widest for conformations with little or no helical structure and gradually narrowed as the number of residues in the helical state increased. Multiple local minima were observed within this landscape, corresponding to conformations with different integer numbers of residues adopting helical structures. The deepest minimum corresponded to the fully disordered state. Repeat 3 showed a somewhat different distribution compared with the other repeats along the radius of gyration and S_helix_ coordinates. This difference may be attributed to limited sampling of conformations with S_helix_ > 10. Nevertheless, the overall shape of the conformational landscape was consistent across all four repeats.

Similar behaviour was observed in the OneOPES simulations. FESs obtained from the conventional MD ensembles or reweighted ensembles of the OneOPES simulations are almost identical. This demonstrates more efficient conformational sampling of FCP1_term_ with OneOPES. In the enhanced sampling simulations, we can again see the crucial role of Lys951 residue in the stable helix formation. A detailed analysis of the OneOPES simulations is provided in the Supplementary Results 1, Figure S2.

To assess the relevance of these conformations to the experimental ensemble, we calculated NMR chemical shifts from the simulation trajectories (Supplementary results 2; Figures S4–S6). Conventional MD and reweighted OneOPES produced similar chemical-shift profiles, indicating that the observed conformational populations are not sampling-limited. In contrast, the well-tempered ensemble showed improved agreement with experiment, suggesting a larger contribution of rapidly interconverting helical conformations. Together, these results indicate that transiently structured states contribute substantially to the equilibrium ensemble of FCP1term.

In all simulations, FCP1_term_ generally followed the same helix-formation mechanism. Short helical structures preferentially form in the 942–951 and 954–960 regions, while Lys951 incorporation is associated with the formation of the extended helical structure. These observations provide an example of a pre-structured motif (PreSMO) in action, suggesting that FCP1term can transiently populate helical, binding-competent motifs in solution before interacting with Rap74. Thus, although the fully extended helix is not the most stable conformation in solution, FCP1_term_ can sample this binding-competent conformation for hundreds of nanoseconds.

### Topological analysis revealed FCP1_term_ folding mechanism

Next, we analysed the conventional MD trajectory using circuit topology which provides a contact-based description of intramolecular organization and enables us to characterize folding transitions (Golovnev & Mashaghi, 2020; Hammond et al., 2026; Heidari et al., 2020). First, we plotted the number of parallel, series, and cross arrangements of contacts as a function of time (Figure 2A). We observed very few topological arrangements, consistent with the disordered state of FCP1_term_, which forms very few intrachain contacts. However, several events increased the number of series and cross-arrangements. The three largest events are highlighted with red rectangles. These events correspond to the formation of the helical structure, as the helix is enriched in cross and series and almost completely lacks parallel arrangements. In contrast, the beta sheet structure is typically enriched in parallel arrangements, with a different pattern of cross arrangements. The distribution of these topological arrangements along the chain for the different structures is shown in Figure S3A. Beta sheet formation events can be identified by peaks in the number of parallel arrangements (Figure 2A). The two largest events are highlighted with blue rectangles.

**Figure 2.**
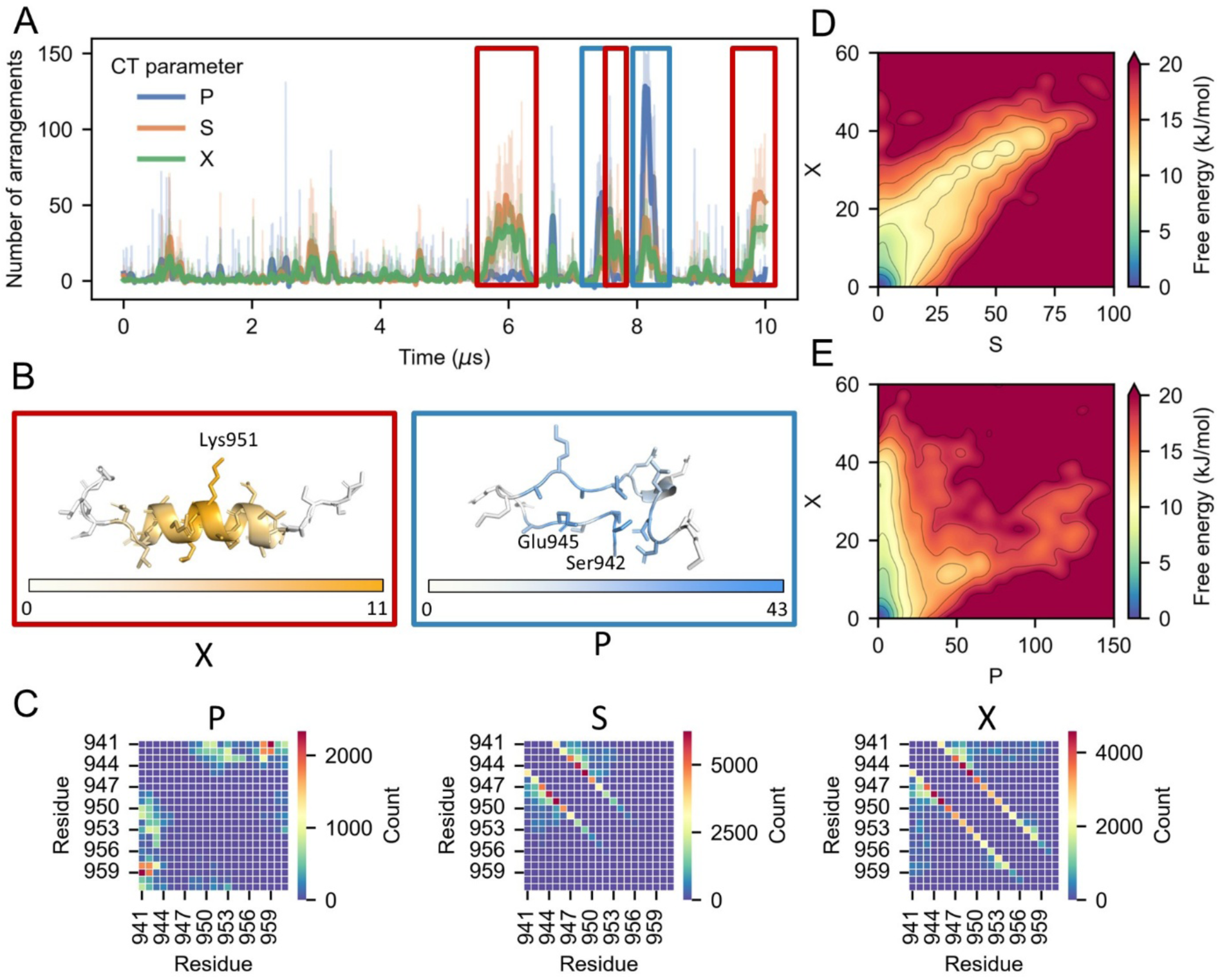
Characterization of the FCP1term folding with circuit topology. (**A**) Dynamics of parallel (P), serial (S), and cross (X) topological contact arrangements during the conventional FCP1term simulation. (**B**) Representative helical (left) and beta sheet (right) conformations, colored by the number of cross and parallel arrangements per residue, respectively. (**C**) Color maps showing, from left to right, the total number of parallel, series and cross arrangements per contact. (**D**) 2D FES projected onto the number of series (S) and cross (X) arrangements. (**E**) 2D FES projected onto the number of parallel (P) and cross (X) arrangements.

In Figure 2B, we show representative structures of the helix and beta sheet together with their corresponding topological arrangements. In the helical structure, Lys951 is involved in the largest number of cross arrangements. The number of cross contacts decreases from this central residue towards both sides of the peptide. In the beta sheet structure, the distribution of parallel arrangements has a different pattern. Here, the terminal residues form the most contacts. In this structure, Ser942 and Glu945 form the most parallel contacts.

Different residue pairs have distinct propensities for each type of topological arrangement. Most parallel contacts are formed by residues near the N- and C-termini. (Figure 2C). Series arrangements are mainly associated with local contacts, particularly between residues 941–949, which are close to the residues that form most of the helical structure (Figure 1C). In contrast, cross arrangements involve local contacts distributed across the entire peptide chain. Interestingly, the patterns for CP and CP^-1^ are different from those observed for P arrangements. They include both local and terminal contacts (Figure S3B), while the pattern for CS is very similar to that of S arrangements. Importantly, series and cross arrangements are the most frequent topological arrangements in FCP1_term_, which may be characteristic of helix-forming IDPs (Hammond et al., 2026).

The 2D FES shows how helical and beta sheet structures are distributed in topological space. The most stable state is the disordered state, characterized by a low number of topological arrangements. Based on the number of cross and series arrangements, the fully formed helix, characterised by approximately 50 series arrangements, is about 10 kJ/mol less stable than the completely disordered state (Figure 2D). A similar analysis can be performed for the beta sheet structure. The region with approximately 120 parallel arrangements corresponds to the beta sheet structure in Figure 2E. This state is around 15 kJ/mol less stable than the disordered state.

Interestingly, the helical structure connects to the disordered state through many intermediate states. This continuous distribution of intermediates is consistent with a gradual formation of the helix as a one-dimensional object and can be interpreted within the framework of phase transitions in one-dimensional systems (L.D. Landau and E.M. Lifshitz, 1980). In this picture, the progressive formation of topological contacts resembles a continuous, second-order-like transition, in which structural order develops gradually through intermediate states. In contrast, the beta sheet structure appears as a much more isolated state in the topological space, suggesting a different mechanism of structural formation.

Thus, circuit topology provides a complementary way to characterize the formation of both helical and beta sheet structures in FCP1term and allows us to distinguish their different structural pathways.

### The FCP1_term_:Rap74 complex retains a dynamic binding interface

We performed conventional MD simulations of the FCP1term and Rap74 complex. The initial state was taken from the 1onv PDB structure. We simulated the system for 10 µs at 300 K and 350 K. We observed that, despite the complex not being disrupted during the modelling time, there were multiple rearrangements in the Rap74 structure, the FCP1_term_ helix structure, and their relative orientations.

Rap74 remains well folded in *holo* state, maintaining three α-helices (H1–H3) and an antiparallel β-sheet, with the main flexibility occurring in the loop between H2 and H3 (Figure 3A) (Supplementary Results 3). Hydrophobic and charged interactions stabilize the Rap74 complex (Figure 3B,C), in agreement with previous reports here (Kamada et al., 2003; Nguyen, Abbott, et al., 2003; Wostenberg et al., 2011). tICA analysis showed that conformational fluctuations are concentrated in the loop and N-terminal disordered region of the Rap74, revealing four distinct conformational clusters (Figure S7A–C). Temperature-dependent analysis showed that Rap74 remains mainly in states 1–2 at 300 K, whereas higher temperature (350 K) promotes more frequent transitions and partial loss of helical structure (Figure S7D,E). Overall, increased temperature primarily induces local structural changes while the global Rap74 fold remains preserved (Figure S7E).

**Figure 3.**
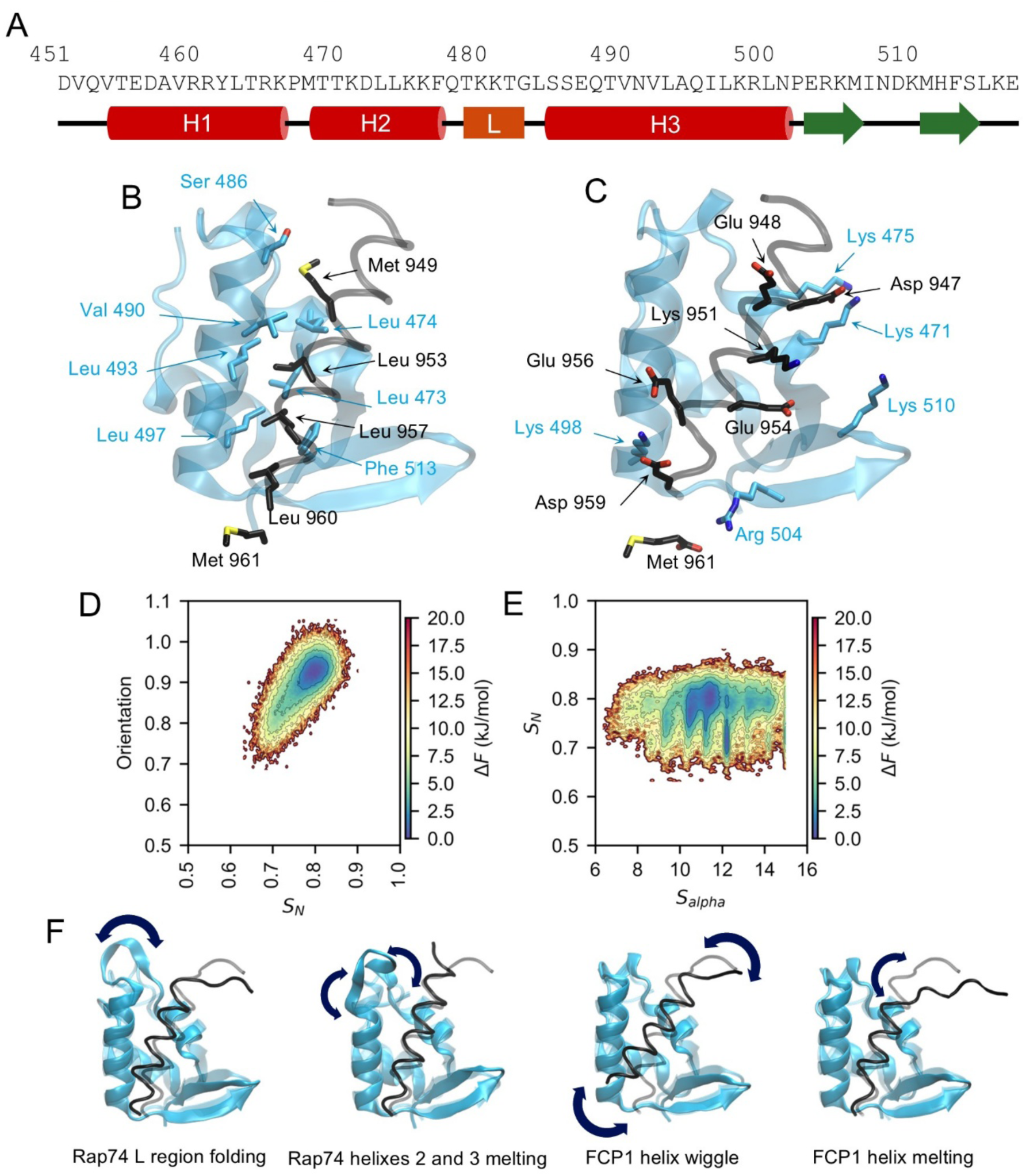
Dynamics of the FCP1term:Rap74 complex. (**A**) Amino acid sequence and secondary structure of Rap74. (**B**) Hydrophobic interactions between Rap74 and FCP1term that contribute to complex stabilization. (**C**) Same as (B), but for charged interactions. (**D**) FES projected onto (SN) and helix orientation. (**E**) FES projected onto helicity and (SN). (**F**) Representative structural ffuctuations observed in the native Rap74:FCP1term complex.

During conventional simulations at 300 K, we observed the fuzzy nature of FCP1_term_ binding to Rap74. In particular, the folded FCP1_term_ helix underwent substantial reorientation within the binding pocket. To characterize this motion, we introduced an orientation CV based on the normalized projection of two vectors: one describing the direction of the binding pocket and the other describing the orientation of the FCP1_term_ helix (Figure S8A). The pocket vector connects the Cα atom of Pro502 with the & atom of Leu474, while the helix vector connects the Cα atom of Glu956 with the N atom of Met949. The orientation CV ranges from −1 for antiparallel vectors to +1 for parallel vectors. The FCP1term helix sampled a range of orientations within the binding pocket, with deviations from the parallel orientation associated with an increase in free energy (Figure 3D).

The FCP1_term_ helix also exhibited substantial conformational fluctuations during the simulations at 300 K. We characterized these fluctuations using the S*_helix_* and S*_N_* CVs, where S*_helix_* describes the number of helical turns in FCP1_term_ and S*_N_* describes the number of native contacts between FCP1_term_ and Rap74. The resulting FES (Figure 3E) shows that the number of native contacts remained relatively stable, fluctuating between 0.7 and 0.9, whereas S*_helix_* exhibited substantially larger fluctuations. The lowest-free-energy region was centered at S*_helix_* = 10–12, but the peptide occasionally sampled states with S*_helix_* ≈ 6 and extended helical conformations with S*_helix_* ≈ 15. Thus, the FCP1_term_ helix can undergo substantial structural fluctuations while maintaining a relatively stable network of native contacts with Rap74.

Additionally, we applied tICA to characterise FCP1_term_ dynamics within the Rap74 binding pocket. First, we considered only the native interchain contacts observed in the complex structure. Using the 1ONV PDB structure, these contacts were defined according to equation 1, with residue pairs considered to be in contact when the resulting value was below 0.75. We then projected the resulting contact space onto the first two tICA components using a lag time of 50 ps. The selected residue pairs and their corresponding tICA weights are shown in Figure S8B,C.

Projection of the trajectories onto the first two tICA components revealed four distinct clusters representing microstates (Figure S8D). At 300 K, the system predominantly occupies states 1 and 2, whereas increasing the temperature promotes transitions to the higher-energy states 3 and 4.

The temperature dependence is also evident in the time evolution of the tICA components, with higher temperature leading to more frequent transitions between the identified states (Figure S8E,F).

Unlike for the Rap74 analysis, both tICA components were required to describe the conformational dynamics, as they captured distinct structural processes. Transitions between states 1 and 2 and between states 3 and 4 along tICA2 correspond to changes in the orientation or “wiggling” of the FCP1term helix within the binding pocket. In contrast, tICA1 primarily captures changes in interatomic distances associated with partial melting of the loop region L of Rap74. The corresponding structural changes are illustrated in Figure S8G.

Despite these conformational fluctuations, the core of the FCP1term binding interface remains relatively stable. Residues directly contacting Rap74, particularly Leu953 and Leu957, remain predominantly helical and maintain hydrophobic interactions with Val490, Leu493, Leu474, Leu473, and Phe513 of Rap74. Leu953 and Leu957 therefore appear to anchor the peptide while allowing the remainder of FCP1_term_ to undergo substantial structural rearrangement. The result is a dynamic complex in which a relatively stable hydrophobic interaction core is embedded within a highly flexible conformational ensemble.

We also analysed the stability of FCP1_term_ helical structure. We aligned all trajectories to the Rap74 structure and calculated the root-mean-square fluctuations (RMSF) of the Cα atoms of FCP1_term_. As shown in Figure S8H, residues directly involved in contacts with Rap74 exhibited the lowest fluctuations, whereas the N- and C-terminal regions of FCP1_term_ showed substantially greater flexibility. In these terminal regions, the helical structure could transiently or completely melt, even at 300 K.

Together, these observations support a fuzzy-complex model for the Rap74:FCP1term complex. Despite substantial conformational fluctuations, including changes in helix orientation and conformation changes of the peptide and Rap74 binding region, FCP1term remains associated with Rap74, and the complex’s overall integrity is preserved. All detected structural changes are listed and illustrated in Figure 3F.

### Enhanced sampling reveals a multimodal folding-and-binding landscape of Rap74:FCP1_term_

To investigate the folding and binding of FCP1_term_ to Rap74, we performed an enhanced sampling simulation of this system. Before processing the OneOPES simulation, we investigated the free energy surface with exploratory simulations to identify a probable metastable state. We used the OPES_METAD_EXPLORE simulation protocol with two collective variables: LDA_interchain_ – native contacts between two protein chains; and LDA_helix_ – backbone hydrogen bonds forming an alpha helix in FCP1_term_. All contacts were weighted with coefficients determined from a short preliminary simulation (see details in the Methods section). With these two CVs, we aimed to increase sampling of folding-upon-binding events in our system. To limit sampling of the medium, we applied a funnel restraint (Limongelli et al., 2013). We performed four independent runs, each lasting 8 µs.

During the simulations, we observed a few unbinding and binding events and several metastable states (Figure 4A). The first metastable state (N) is close to the canonical known folded structure with FCP1term in the helical conformation, forming native hydrophobic and charged contacts. In the second metastable state (F), FCP1_term_ also forms helix, but this helix is flipped relative to the native orientation. This state was also stabilized by hydrophobic and charge contacts. Because FCP1_term_ has some level of symmetry, flipped and folded peptides can form contacts with the same residues of Rap74 as in the native conformation (Figure S9). The disordered-bound conformation (D) is an ensemble of conformations in which mainly native hydrophobic contacts form, but FCP1_term_ remains disordered. All these states could be seen in the free energy surfaces (Figure 4B).

**Figure 4.**
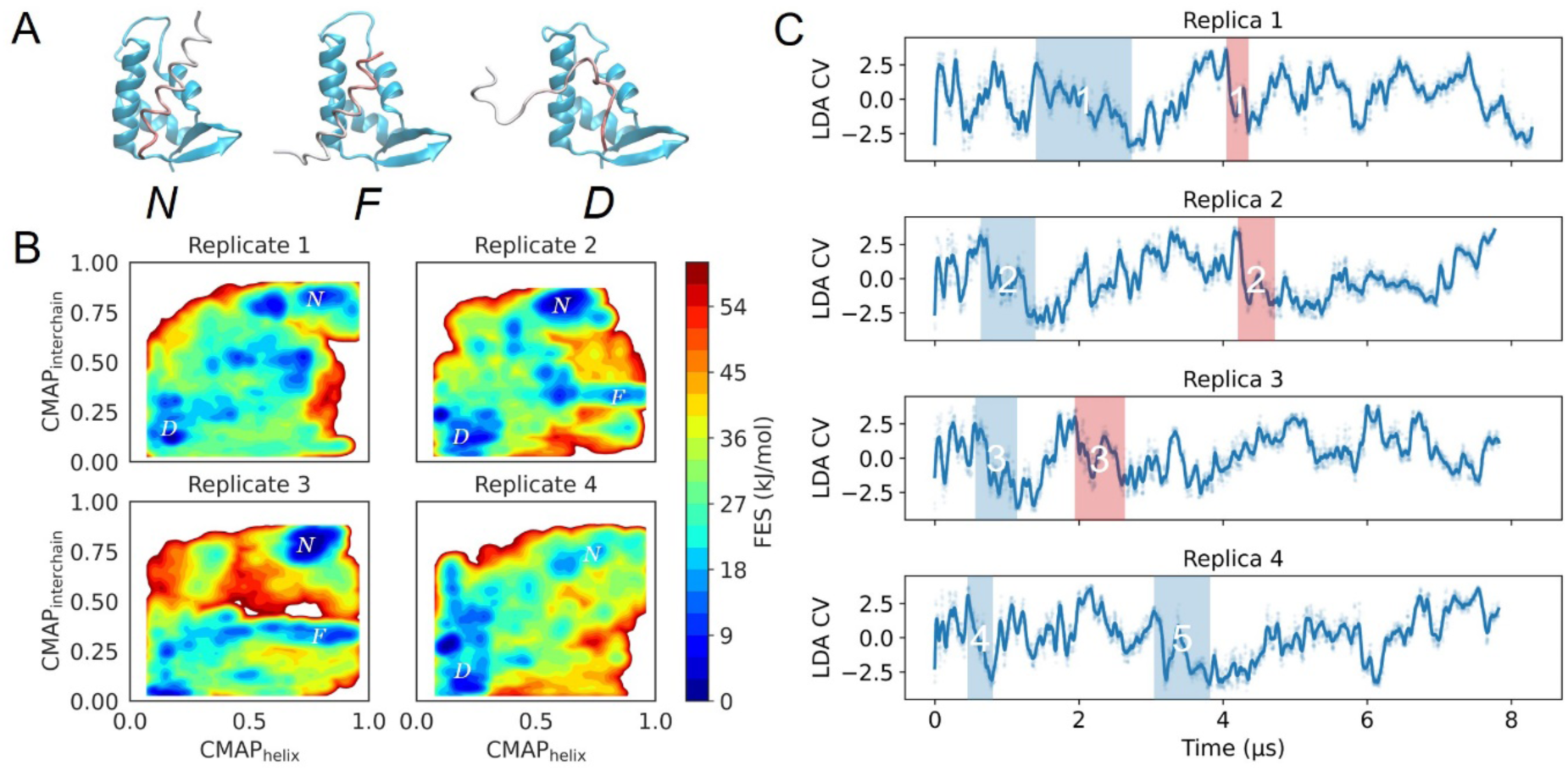
Multimodal binding of Rap74:FCP1term complex. (A) Representative structures of the native complex N, ffipped complex F and disordered complex D. FCP1term is colored with a gradient from white to red, from the N-terminus to the C-terminus. (B) Reweighted free energy surface from biased simulations for each replicate. CMAPinterchain is the normalized sum of native interchain contacts between proteins. CMAPhelix is the normalised sum of CO - HN atom contacts that form hydrogen bonds in the alpha-helical structure of FCP1term. Significant deviations in the height of free energy surfaces are observed between different runs; however, in all simulations, a correctly folded and bound (F) basin is observed. In replicates 2 and 3, basins of ffipped and bound states (F) are observed. The disordered bound (D) state populates the lower-left part of the plots. (C) Dynamics of the protein complex in OPES Explore biased simulations traced with the LDA CV. Five folding events were observed. They are marked with blue boxes. We also observed three misfolding events, highlighted with red boxes.

The flipped state is stabilised by non-native hydrophobic and charge contacts between FCP1_term_ and Rap74 (Figure S9). Compared with the native state, in the flipped state Leu 953 remains in place, whereas Leu 957 and Met 949 change. Interestingly, new interactions between Glu 956 and Lys 471, and between Arg504 and Glu945 and Glu948, are formed.

Despite aggressive sampling, the free energy surfaces obtained in this set of experiments did not converge. This is clearly seen in the significant differences between the FESs for each repeat (Figure 4B). Some features of the FES look similar, like the location of the N or F states, but not in all repeats. For example, the stable native state was detected in replicates 1, 2, and 3. The flipped state was only observed in replicates 2 and 3. Dynamics in the CV space do not show sharp transitions from one state to another, but rather a slow diffusional process. (Figure 4C). This could be explained by imperfections in the CVs, when different conformations occupy the same regions of CV space. For example, this CV does not distinguish the F state from intermediates towards the natively folded state. If the flipped state is metastable, the system gets trapped in that state until the bias can adapt. Nevertheless, we identified important intermediate metastable states.

“Divide et impera”: thermodynamic decomposition enables absolute binding free-energy calculation

We investigated the folding and binding process using OneOPES simulation. As shown with conventional MD, a 10 µs simulation is not sufficient to fully sample unbinding. With the help of the OPES biasing scheme, we not only observed several unbinding and rebinding events but also produced a converged FES. This information helped us propose a model of the folding and binding of FCP1term to Rap74.

The OPES method uses the CVs, along which the algorithm aims to build a converged FES. From preliminary simulations, we observed that accounting for only native contacts is insufficient to obtain converged data. A good CV should not only distinguish the initial and final states but also the transition state. In the preliminary simulations, we observed the formation of a flipped state, which had not been previously described. When this metastable state occupied a region similar to the transition states along the path from the native complex to the apo state, the simulation became trapped in it. Interestingly, this state was populated because FCP1_term_ has some symmetry. If the helix were to flip, the hydrophobic and similarly charged residues would remain in similar positions.

### Thermodynamic cycle

The free energy of FCP1_term_ folding and binding to Rap74 Δ*F_folding-binding_* was estimated using the thermodynamic cycle shown in Figure 5A, which separates the overall process into peptide folding and binding of the folded peptide. The folding free energy Δ*F_folding_* was calculated for isolated FCP1_term_, based on the assumption that, in the unbound state, Rap74 does not significantly influence the conformational ensemble of the peptide. And at the same time free energy of solubilization Δ*F_Rap74_* of *apo* Rap74, does not depend on the folding of FCP1_term_.

**Figure 5.**
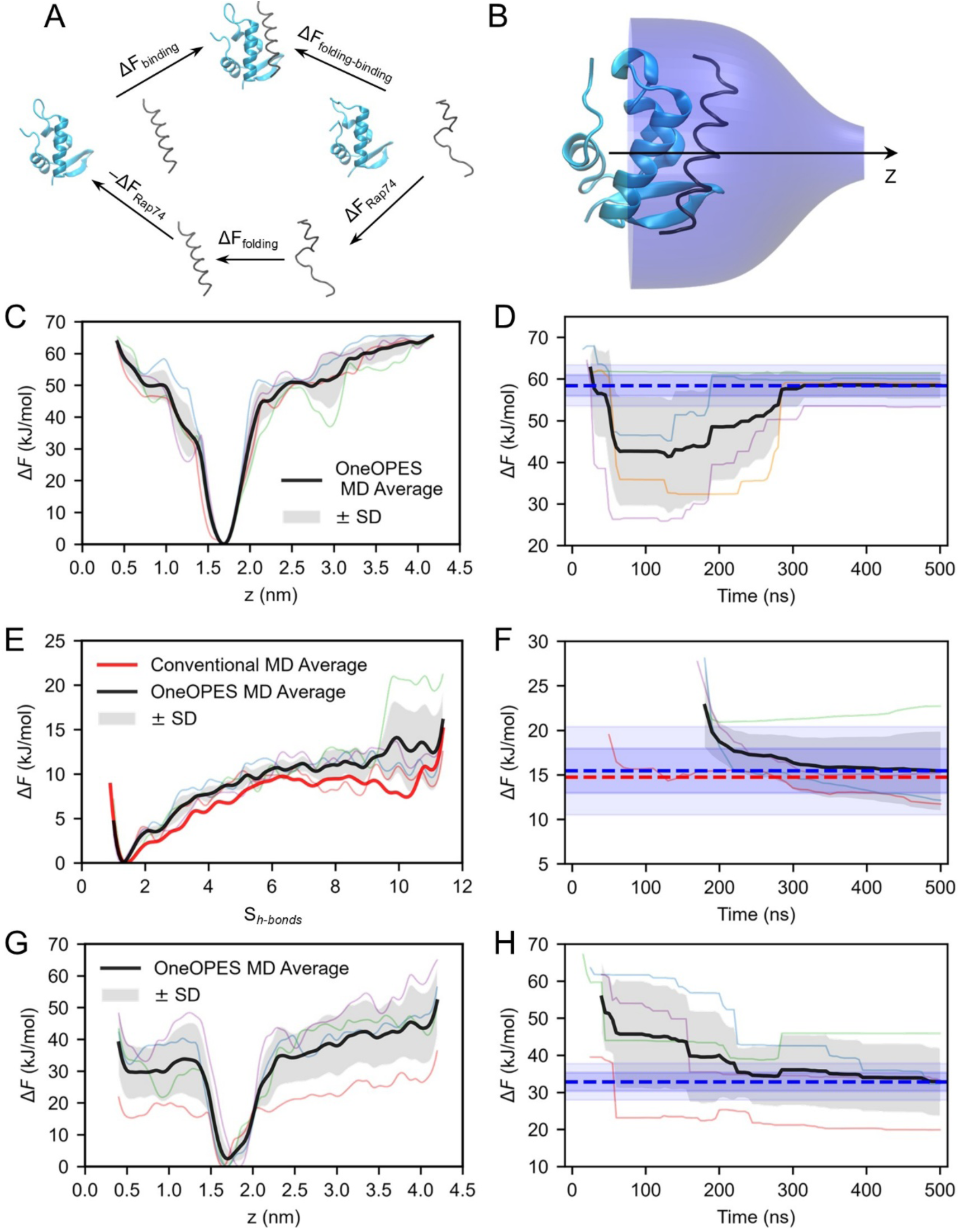
Free energy calculation of coupled folding and binding. (**A**) Thermodynamic cycle connecting the folded and bound state with the free and disordered state. ΔFfolding-binding was calculated from the free energy of binding of the peptide in its helical state, ΔFbinding, and the free energy of folding of the isolated peptide, ΔFfolding. In the unbound state, we assume that the conformation of FCP1term does not affect Rap74’s conformation; therefore, we subtract the free energy associated with Rap74 solubilization, ΔFRap74, within the thermodynamic cycle. (**B**) Simulation system showing Rap74 in cyan, FCP1term in black, and the funnel restraints in blue. The black arrow indicates the symmetry axis of the funnel restraints. (**C**) Free-energy surface of the centre of mass of FCP1term in the helical state projected along the z-axis. Coloured lines represent the FESs from independent repeats; the bold black line represents their average, and the shaded area represents the standard deviation. (**D**) Convergence of the free energy of binding. The notation is the same as in (**C**). The horizontal blue dashed line indicates the converged average free energy of unbinding, 58 ± 3 kJ/mol. The dark and light blue shaded areas represent regions within 1 kBT and 2 kBT of the converged value. (**E**) Same as (**C**), but with the FES projected along the Sh-bonds collective variable. The bold red line represents the FES obtained from conventional MD simulations. (**F**) Convergence of the free energy of folding, ΔFfolding. The notation is the same as in (**D**). The bold blue dashed line is the converged free energy of folding, 15 ± 4 kJ/mol, calculated from the OneOPES simulations. The bold red dashed line represents the free energy of folding calculated from conventional MD, 14.7 kJ/mol. (**G**) Free-energy surface of the centre of mass of FCP1term projected along the z-axis. The notation is the same as in (**C**). (**H**) Convergence of the free energy of folding and binding ΔFfolding-binding. The notation is the same as in (**G**). The horizontal blue dashed line indicates the converged average free energy of unbinding, 33 ± S kJ/mol.

The binding free energy Δ*F_binding_* was calculated for the FCP1_term_ in the helical state. The helical conformation of the peptide region involved in the native and flipped bound states was maintained by restraining the corresponding intrachain hydrogen bonds. As these restraints were applied only to the binding-relevant region, the rest of the peptide could adapt disordered conformation. By limiting sampling of fully disordered conformations, this approach reduced the conformational space explored during the binding simulations while retaining the conformations relevant to the binding mechanism. This restraint still allows us to estimate the free energy difference between N and F states.

The separate folding and binding free-energy contributions can therefore be combined to estimate the overall free energy of the binding process.

### Free energy of binding

Taking into account F state we investigated the process of FCP1_term_ binding to Rap74. We performed the OneOPES simulation of FCP1_term_ to Rap74 using a new set of CVs: *S_N_* – normalized sum of the interchain contacts within N state, S_F_ – normalized sum of the interchain contacts within F state, and z – the distance between center of mass of CA atoms and binding pocket. Additionally, we used one supporting CV, the radial distance from the center of mass of the ligand to axis z. This time, all contacts, had an equal weight. Same as in the previous case, we used a funnel restraint (Figure 5B).

Including the non-native contacts characteristic of the F state facilitated the identification of the transition region between the apo, N, and F states. The resulting three-dimensional FES has a characteristic Y-shaped topology, with the two upper branches leading toward the N and F bound states and the lower branch leading toward the apo state. The vertex of the Y-shaped landscape corresponds to a region with few native or flipped contacts, providing a distinct representation of the transition between the three states. Four independent simulations were performed, each comprising eight OneOPES replicas simulated for 500 ns, corresponding to a total of 16 μs of sampling. Further details of the simulation setup are provided in the Methods section.

Using OneOPES simulation protocol, we calculated the FES along vector ***z*** (Figure 5C). The final FES profiles are quite similar across all four repeats. The minimum of the surface is located around the initial state, and after a rapid increase, it approaches a plateau. This profile shows that the bound state is the most stable one. We used a cutoff of ***z*** > 3.5 nm, to define the free state, as a helical peptide can still make contacts with Rap74, when it is positioned closer. This could be nicely seen, as in one of the repeats (green line) we had a local minimum around 3 nm distance, when a helical peptide was oriented along the vector ***z***. Then we calculated the free energy difference in the reweighted ensemble. Here we calculated the free energy of unbinding Δ*F_un_*_binding_, where Δ*F_un_*_binding_ = –Δ*F_binding_*. All simulations converged at around 300 ns of the simulations (Figure 5D). We calculated the average free energy of binding as -58 ± 3 kJ/mol. Importantly the uncertainty is less than *2*k_B_T ≈ 5 kJ/mol. However, this value is not final, as we need to take into account the correction, representing effect of the funnel restraints, that was -10 kJ/mol. Then we receive Δ*F_binding_* = -48± 3 kJ/mol.

Each replicate sampled the relevant conformational space extensively, as shown in Figure S10A–D. For clarity, only the data from the first replica are presented in these plots. State N (*z* ≈ 1.7 nm, *S*_N_> ≈ 0.8) accumulated the largest bias. Importantly, this state was revisited at least once after peptide unbinding, which was necessary to achieve convergence and obtain an accurate estimate of the bias. Interestingly, although state F was also visited multiple times, it consistently accumulated a lower bias than state N. This indicates that state F is less stable than the native state.

Figure S10E shows two-dimensional free-energy surfaces, where *z* represents the projection of the peptide centre of mass onto the *z*-axis, and *R* represents the radial distance of the peptide centre of mass from the same axis. All four replicates show similar free-energy landscapes, with the peptide-bound state representing the most stable region. Importantly, the funnel restraint did not introduce new metastable states, and provided an efficient connection between protein-bound and peptide-free states.

Figure S10F compares the relative stability of the native S_N_ and flipped S_F_ states. The native state is more than 30 kJ/mol more stable than the flipped state, confirming that state F represents a substantially less stable, non-native binding configuration. Despite its lower stability, sampling of the flipped state was important for the simulations. The introduction of these non-native contacts helped define the transition region between the bound and unbound states and contributed to the construction of an efficient CV, thereby facilitating convergence of the simulations.

### Free energy of folding

We next used the OneOPES folding simulations to estimate the free energy of folding, Δ*F_folding_*. For this analysis, we introduced an additional collective variable, S*_h-bonds_*, which describes the number of hydrogen bonds formed that were restrained during the binding simulations. A fully folded structure can form a total of 11 such hydrogen bonds.

Figure 5E compares the free-energy surfaces obtained from conventional MD and OneOPES simulations. The OneOPES free-energy surfaces were calculated using the reweighted ensemble. Overall, the free-energy profiles obtained from the two simulation approaches were similar. The energy minimum is around 0, indicating disordered nature of the peptide. Increasing the number of hydrogen bonds increases the free energy of the system. Three of the four OneOPES repeats showed consistent profiles, whereas repeat 3 (green) differed at high values of S*_h-bonds_* > 9. This was the same repeat that showed differences in the conformational landscape discussed in the previous section.

The free energy of folding was subsequently estimated from these free-energy surfaces. The folded state was defined based on the formation of the hydrogen bonds described by S*_h-bonds_*. The convergence analysis showed that, after approximately 300 ns, Δ*F_folding_* fluctuated within approximately one *k*_B_*T*, indicating convergence of the free-energy estimate.

### Final free energy difference

The converged average folding free energy of folding Δ*F_folding_* obtained from the OneOPES simulations was 15 ± 4 kJ/mol (blue dashed line), in close agreement with the value obtained from conventional MD (14.7 kJ/mol, red dashed line), despite the different behaviour observed for repeat 3. Excluding repeat 3 would result in a lower estimate of the folding free energy from the OneOPES simulations.

Finally, we calculated the absolute free energy of folding and binding as: Δ*F_folding-binding_ =* Δ*F_binding_ +* Δ*F_folding_ = –*33 ± 5 kJ/mol. This value is close to the experimentally determined –32.5 ± 0.5 kJ/mol (Prieto et al., 2021) or –36.2 ± 0.5 kJ/mol (Yang et al., 2009). Thus, our calculation and proposed method quantitatively reproduce the experimental free energy of folding and binding for the disordered peptide.

### Direct estimation of the folding and binding free energy

To estimate the efficiency of the simulation *Divide et impera* approuch, we calculated the free energy of folding and binding with the OneOPES protocol. We used the same CVs, and funnel restraint as used for Δ*F_binding_* calculations, but added the number of helical segments in FCP1_term_ S*_helix_* to promote dynamics of helix rearrangements as one more supporting CV.

The free energy surfaces of the centre of mass of the FCP1term along axis z feature a minimum around 1.7 nm and a plateau at 3.5 nm (Figure 5G). In contrast to Figure 5C, the deviations between repeats are much more pronounced. Several repeats show plateaus near 40 kJ/mol, while one plateaus near 25 kJ/mol (red line). The average free energy of unbound FCP1_term_ is around 60 kJ/mol for the restricted helix case, whereas it is around 40 kJ/mol here. This difference could be attributed to FCP1_term_’s greater flexibility in the non-restricted helix, where the peptide structure could adapt to the solvent.

The standard deviation in the convergence plot (Figure 5H) exceeds 2 kBT, despite low fluctuations within each individual repeat, indicating that the sampled conformational space is not identical. The average of the converged free energy of binding is –33 ± 9 kJ/mol. After applying the funnel-restraint correction, we get ΔF_folding-binding_ of –23 ± 9 kJ/mol. This value is close to the largest experimental estimate; however, its uncertainty is larger than that calculated from a thermodynamic cycle.

It is interesting to analyse why we received a different FES for each of the repeats. As we can see in Figure S11A, all repeats show well-populated bound and free states, with data points well distributed along the z-axis. However, only in repeats 2-4 the FCP1_term_ revisited the natively bound (Figure S11B) and partially folded (Figure S11C) structures, as can be seen from the data distribution and the amount of bias. At the end of repeat 1, FCP1_term_ formed native contacts but lacked a helical conformation, resulting in a high-free-energy state. Such differences in sampling resulted in different FESs (Figure S11D-E). Again, repeats that managed to revisit a low-energy N state have a convex shape.

From the free-energy surfaces, shown in Figure S11E, the native state is more stable than the flipped state, consistent with the simulations with restricted helical conformations. However, the free-energy difference between these states varies substantially between replicates, ranging from approximately 10 to 40 kJ/mol.

Finally, we analysed the changes in the topological organisation in the presence of Rap74. To do so, we built a reweighted FES projected onto the cross-parallel P and cross-X axes and the series and cross topological arrangement axes (Figure S12A, B). In contrast to Figure 2D, E the most stable conformation has X>20 and P<20. This range is typical for helical structure. We interpret this as the helix stabilisation on the surface of Rap74. Importantly, in contrast to solvent exposed case, we only observed formation and stabilisation of parallel topology in repeat 2. Thus, CT confirms that FCP1_term_ peptide complex topology is stabilised by the Rap74.

### Mechanism of folding-and-binding: distinct pathways converge on the native complex

OneOPES simulations reveal that FCP1_term_ can reach the native Rap74-bound state through multiple pathways, including binding of a partially folded peptide followed by progressive helix formation and native-contact consolidation (Figure 6A–D, **Supplementary Results 4**). Interestingly, the peptide can also form a disordered but partially bound complex, suggesting that nonspecific hydrophobic interactions can stabilize encounter states that subsequently evolve toward the native complex. Analysis of native-contact formation indicates that interactions involving Leu953 and Leu957 form at the earliest stages of binding, followed by contacts involving Met949 and then the remaining native interactions. Helix formation overlaps with this process: a short helical segment involving the central region of FCP1_term_ emerges before the complete helix is established, supporting a coupled rather than strictly sequential folding-and-binding mechanism. More information on the folding-to-binding mechanism is provided in **Supplementary Results 4**.

**Figure 6.**
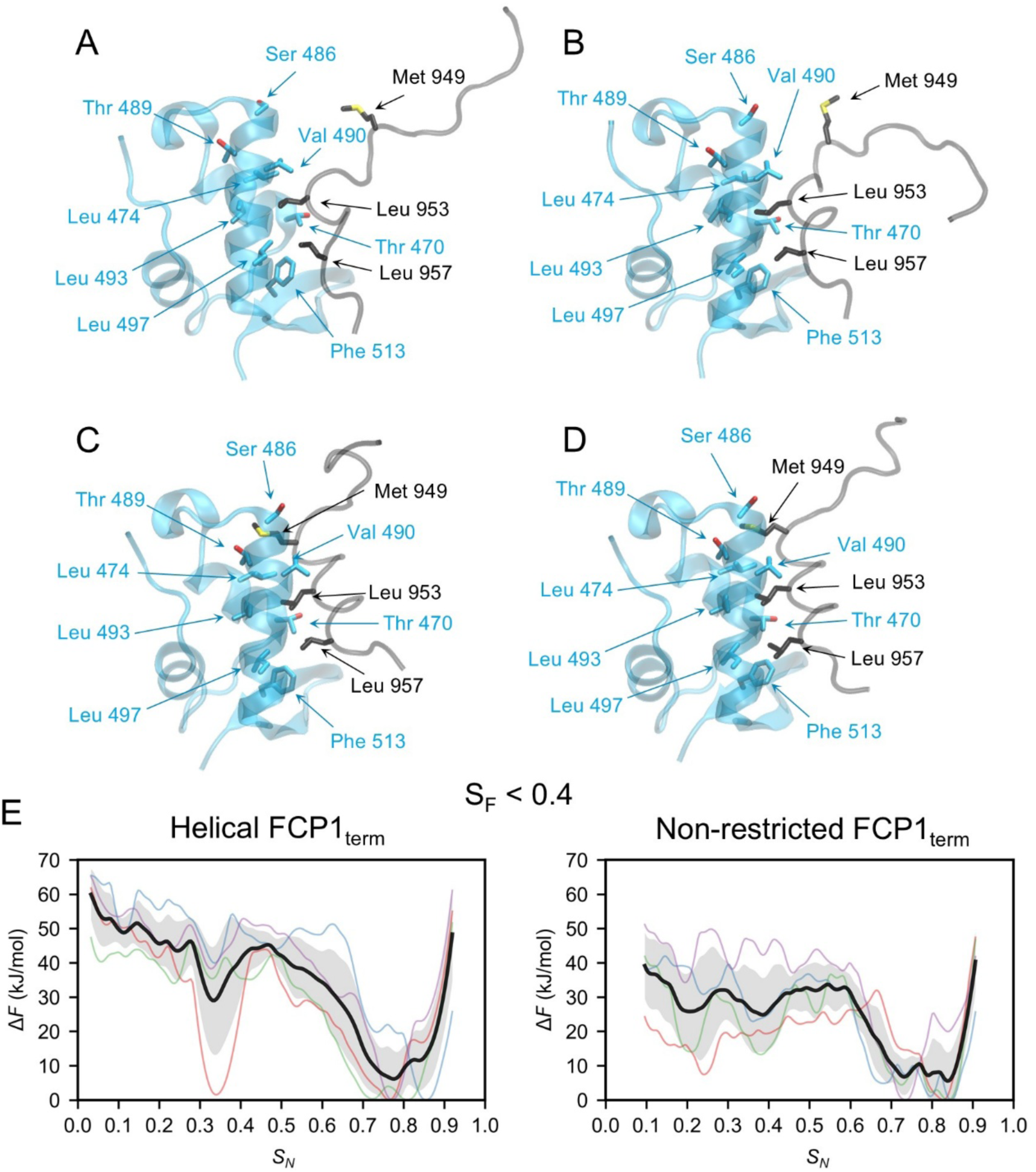
Coupled folding and binding of FCP1term to Rap74 via induced fit mechanism. (**A**-**D**) Representative frames illustrate a binding route consistent with the induced fit. Key interacting residues are shown and labelled. (**E**) Comparison of the FES projected onto SN CV, for SF < 0.4. These profiles represent the change in the free energy along binding to native state for the helical FCP1term and non-restricted FCP1term OneOPES simulations. The bold black line represents the average FES, and the shaded area represents the standard deviation between repeats.

Alternatively, a pre-folded helical peptide can bind and reorient within the Rap74 pocket before reaching the native configuration. Similar reorientation is also observed in conventional simulations, where changes in helix orientation are associated with transitions between bound and unbound configurations (Figure 3D). Comparison of the free-energy profiles further indicates that the pre-folded peptide progresses more directly toward the native state, whereas the unrestricted peptide samples a broader ensemble of partially bound and partially folded conformations before native-state consolidation (Figure 6E). Together, these results indicate that folding-and-binding can proceed through distinct routes that converge on the same native complex, with the balance between binding and folding determined by the conformational state of FCP1term at the time of association. Detailed free-energy surfaces, contact-formation probabilities, and alternative pathways are provided in Figures S13–S14 and Supplementary Movies 1–2.

## Discussion

In this work, we used enhanced-sampling molecular dynamics, Circuit Topology, and thermodynamic decomposition to characterize the conformational landscape, binding mechanism, and energetics of the intrinsically disordered FCP1term–Rap74 complex. In solution, FCP1term predominantly samples disordered conformations but can transiently form extended α-helical and β-sheet structures, as shown with circuit topology. We identified Lys951 as a key determinant of extended helix formation. In complex with Rap74, FCP1term retains substantial conformational flexibility, consistent with a fuzzy complex. Leu953 and Leu957 form persistent hydrophobic contacts that anchor the peptide, while the helix can reorient and its terminal regions can reversibly unfold. Our approach revealed an alternative flipped state and enabled characterization of the coupled folding and binding process. Comparison of the simulated pathways suggests that both conformational selection and induced fit can contribute to complex formation. Early hydrophobic contacts involving Leu953 and Leu957 were observed along both routes.

### Conformational landscape and topological organization of FCP1_term_

Our simulations of FCP1term in solution indicate that the peptide samples both disordered and α-helical conformations. The formation of an extended helix appears to be associated with the incorporation of Lys951 into the helix, which enables a network of intrachain hydrogen bonds involving Glu947 and Glu954, thereby stabilising the longer helical structure. However, the extended helix is only transiently populated. Its formation brings Met and Leu residues into proximity, generating a hydrophobic patch that subsequently interacts with Rap74. This hydrophobic patch may stabilise the helix. The balance between favourable intrachain interactions and the energetic cost of exposing hydrophobic residues to solvent may thus contribute to the conformational equilibrium between the disordered and helical states of FCP1_term_.

This interpretation is consistent with previous work by Gibbs and Showalter (Gibbs & Showalter, 2016), who used Monte Carlo and MD simulations to show that interactions involving Glu947 and Lys951 are important for formation of the FCP1term helix. Together, these observations suggest that FCP1_term_ dynamically exchanges between disordered and partially or fully helical conformations, a behaviour characteristic of PreSMos.

Circuit Topology provides a complementary perspective on FCP1term folding by describing conformational changes in terms of the topology of the contact network. Within a unified CT framework, we identified multiple structural states involving α-helix and β-sheet formation. Helical structures were associated primarily with crossing contact pairs, whereas β-sheet structures were characterised by parallel contact arrangements. Importantly, we observed that Rap74 stabilises helical cross-contact topologies. The low number of parallel rearrangements in the presence of Rap74 suggests that Rap74 does not stabilise β-sheet structures. Thus, the topology of the contact network provides a simple representation of the structural changes occurring during folding, without requiring explicit structural classification.

Interestingly, the progressive formation of the helical structure follows a trajectory along the CT landscape that is approximately aligned with the direction previously associated with increasing topological complexity (Hammond et al., 2026). Thus, FCP1_term_ folding may be described not only in terms of conventional structural order parameters, such as helical content, but also as a progressive reorganisation of its contact topology.

### Comparison of simulated and experimental NMR chemical shifts

The comparison of simulated and experimental NMR chemical shifts indicates that the conventional MD ensemble captures key features of the conformational properties of FCP1_term_, although some discrepancies with the experimental data remain. In particular, the calculated chemical shifts suggest a somewhat more disordered conformational ensemble than indicated by the experimental measurements. Importantly, the experimental chemical shifts largely fall within the range sampled by the conventional MD ensemble, suggesting that conformations compatible with the experimental ensemble are present in the simulations.

A similar trend can be observed when comparing the results of the OneOPES simulations in the reweighted and well-tempered ensembles. The reweighted ensemble reproduces conventional MD results, whereas the well-tempered statistics show better correlation with experimental NMR chemical shifts. The improved agreement with experimental NMR shifts suggests that the experimental ensemble may involve more frequent transitions between disordered and helical conformations than those observed in conventional MD simulations. In addition, folded or partially helical conformations may be populated more frequently than predicted by conventional MD and reweighted OneOPES simulations. From the conventional MD simulations, we can estimate the lifetime of the helical structure to be up to hundreds of ns. During this time, FCP1_term_ could be more likely to bind Rap74.

The observed differences may also arise from differences between the experimental and simulation peptides. The NMR measurements were performed with an FCP1 peptide comprising residues 879–961 or 930–961, whereas our simulations included only the isolated FCP1_term_ fragment (residues 940–961). In our simulations, N- and C-terminal residues were not capped. By altering conformational preferences, or simply by moving the N-terminal charge away from the helix, the additional residues could stabilise the helical conformation of FCP1_term_ through intramolecular interactions. The protonated N-terminus competes with Lys951 for contact with Glu 947, forming a turn structure, and repels Lys951 in the helical state. A similar effect has previously been reported (Ortega-Alarcon et al., 2021; Patapati & Glykos, 2010). The longer peptide has a higher affinity for Rap74 (Prieto et al., 2021). In this research, it was shown that FCP1_term_ has K_d_ = 2.2 µM, whereas the peptide with residues 879−961 has K_d_ = 1.0 µM. This 2 kJ/mol difference is close to the 5 kJ/mol difference that we obtained when comparing conventional MD with the average well-tempered ensemble estimation (Figure S6C). However, additional studies are needed to determine the extent to which the extra residues affect the conformational behaviour of FCP1_term_.

Alternatively, the discrepancy between the experimental data and the conventional MD and reweighted OneOPES simulations may be partly due to limitations of the DES-amber force field (Piana et al., 2020), which may underestimate the stability of helical conformations. Further optimisation of the force-field parameters may therefore improve conventional MD simulations’ ability to reproduce the experimentally observed conformational ensemble of FCP1_term_.

### Insights into native complex structure

Previously, FCP1_term_ was shown to form a fuzzy complex with Rap74, in which the peptide retains substantial conformational flexibility while remaining associated with the binding partner(Wostenberg et al., 2011). For example, in the 250 ns simulations reported in that study, Met949 was observed to transiently detach from Rap74. Our simulations extend this picture by revealing conformational dynamics on both sides of the interface. We identified four major types of structural fluctuations that contribute to the fuzzy nature of the native complex.

First, the loop connecting helices H2 and H3 of Rap74 (loop L) can reversibly adopt a helical conformation, resembling the structure observed in the 1J2X crystal structure (Kamada et al., 2003). Second, at elevated temperatures, the helical regions surrounding loop L can partially and reversibly unfold. Third, the FCP1_term_ helix can reorient within the binding pocket. Such changes in orientation are also apparent when comparing the 1ONV and 1J2X structures: the FCP1_term_ helix adopts a parallel orientation relative to the Rap74 binding pocket in 1ONV, whereas it is rotated in 1J2X (Nguyen, Abbott, et al., 2003). This difference is primarily associated with how deeply Leu957 inserts into the binding pocket. Finally, the N- and C-terminal regions of FCP1term can reversibly lose their helical structure, while the central helix remains associated with Rap74.

Several of the conformational changes observed in our simulations are consistent with previous NMR observations for the Rap74:FCP1term complex (Lawrence & Showalter, 2012). In particular, the flexibility of FCP1_term_ within the binding pocket and the persistence of the interaction despite conformational fluctuations support the previously proposed fuzzy-complex model. The NMR study did not report the partial melting of the Rap74 helices observed at 350 K, which may reflect increased thermal perturbation in our simulations rather than a feature of the complex under physiological conditions. Our results suggest that fuzziness in the Rap74:FCP1_term_ complex arises from coordinated flexibility in both binding partners, while a small set of persistent hydrophobic contacts maintains the overall integrity of the complex.

### Mechanism of folding-and-binding

OPES simulations enabled the study of the mechanism of coupled folding and binding. Using adaptive on-the-fly probability-enhanced sampling, we promoted multiple unbinding and rebinding events. In the exploratory simulations, we observed the formation of the flipped state; however, with the OneOPES framework, we demonstrated that this state is 40 kJ/mol less stable than the native state. Additionally, we observed that FCP1_term_ has a broad ensemble of bound conformations on the Rap74 surface, with varying degrees of disorder.

We observed that hydrophobic contacts play a crucial role in complex formation. Contacts between the Rap74 binding pocket and the hydrophobic residues Leu953 and Leu957 form first on the pathway to the native complex. This confirms the experimental observation that Rap74:FCP1 complex formation is under entropic control (Lawrence et al., 2014). During the protein complex formation, their hydrophobic surfaces collapses. Thus. the association increases solvent entropy through the release of water molecules from hydrophobic surfaces.

Interestingly, these interchain contacts were important in simulations both with and without a restrained helical conformation of FCP1_term_. This suggests that FCP1_term_ does not exclusively follow an induced-fit or a conformational-selection mechanism, but rather that both pathways may contribute to complex formation. Consistent with conformational selection, FCP1term can form partial or complete helical structures in isolation, which can subsequently bind Rap74. Unrestricted OneOPES simulations, however, revealed binding events in which Rap74 contact formations occurred before the helical structure was complete, consistent with an induced-fit mechanism. Together, these simulations support a previously proposed atomistic model of a mixed-binding mechanism for the Rap74:FCP1term complex. (Gibbs & Showalter, 2015; Lawrence et al., 2014).

The different shapes of the FES along the binding coordinate provide qualitative insight into the kinetic differences between binding of pre-folded FCP1_term_ and folding-coupled binding on the Rap74 surface (Figure 6E). For the helix-restrained peptide, the free-energy profile decreases progressively toward the native state, whereas the corresponding profile for the unrestricted peptide remains relatively flat at lower values of S_N_, suggesting that additional conformational rearrangements are required before the native complex is reached. The complex structures show a hierarchy of stability. An increase in the number of native contacts decreases the free energy of the system, making the structure increasingly stable, at S_N_ > 0.6 (Figure 6E). However, these FES profiles should not be interpreted as direct measurements of binding kinetics. Conformational selection requires forming a helical FCP1_term_ before binding, but the timescale of helix formation in solution may limit the rate of this pathway. Thus, although binding of the pre-folded peptide may proceed along a more downhill free-energy profile once the helical conformation forms, the timescale required to populate this conformation may limit the overall kinetics of the conformational-selection pathway.

In the present study, we did not quantitatively determine the relative contributions of the induced-fit and conformational-selection pathways. However, we may compare the binding of the folded helix with folding on the surface. Such a comparison requires a flux-based analysis of the binding processes(Greives & Zhou, 2014; Hammes et al., 2009). OPES flooding simulations could provide a framework for quantifying these contributions in future studies. By comparing the binding kinetics of the helix-restrained and unrestricted peptides, along with the kinetics of helix formation in the isolated peptide, it may be possible to estimate the relative rates of the conformational-selection and induced-fit pathways.

### Free energy calculations using “Divide et impera”

Calculating absolute binding free energies remains challenging for highly flexible molecules, including intrinsically disordered proteins and peptides that undergo coupled folding and binding. In these systems, the large conformational space of the unbound ligand is coupled to its interaction with the receptor, making it difficult to identify and efficiently sample the slow collective motions that govern association and folding. Here, we show that this coupled process can be decomposed into distinct thermodynamic contributions from peptide folding and binding. We calculated the folding free energy for the isolated peptide in solution and evaluated its binding contribution using a structurally restricted peptide ensemble corresponding to the bound state. This decomposition reduces the conformational space, thereby providing a more tractable route to absolute free-energy estimation.

This strategy builds on the thermodynamic framework proposed by Hadži and Lah, in which the free-energy cost of pre-folding an intrinsically disordered α-helical motif was treated as a separate thermodynamic contribution to binding (Hadži & Lah, 2022). We applied this concept to FCP1_term,_ and the resulting folding penalty of 15 ± 4 kJ/mol is close to the value of 13.5 ± 0.9 kJ/mol predicted for FCP1_term_ using the model of Hadži and Lah. The binding contribution of the helical peptide was −48 ± 3 kJ/mol. Although conformational fluctuations of FCP1_term_ within the binding pocket could alter the free-energy difference between bound states, the OneOPES binding calculations explicitly sample these fluctuations and therefore incorporate them into the resulting free energy.

An important implication of this decomposition is that the computational pathway linking two thermodynamic states need not replicate the physical binding pathway. Free energy is a state function, so the equilibrium free-energy difference between the initial and final states is independent of the path between these states. This consideration is particularly important for intrinsically disordered peptides, which may bind via multiple pathways, including induced fit and conformational selection. A direct simulation can sample one or several of these physical pathways, depending on the chosen collective variables and sampling protocol, whereas the thermodynamic cycle used here deliberately separates folding from binding. The resulting pathway therefore need not correspond to the dominant microscopic mechanism of association. Instead, it provides a computationally convenient route for evaluating the same equilibrium free-energy difference.

This decomposition substantially improved the convergence of the absolute binding free energy for FCP1_term_. The combined estimate of ΔF_folding-binding_ = −33 ± 5 kJ/mol is close to the experimental value of −32.5 ± 0.5 kJ/mol, whereas the direct calculation gave ΔF_folding-binding_ = −23 ± 9 kJ/mol. Thus, separating the folding and binding contributions reduced uncertainty and produced a free-energy estimate in closer agreement with experiment. Reducing the dimensionality of the conformational space in the binding calculation substantially improved the efficiency of absolute free-energy calculations for flexible ligands.

The approach nevertheless has several limitations. First, the decomposition requires additional simulations compared with a single direct calculation. For short peptides, however, the folding calculation can be performed in a smaller solvent box and is therefore substantially less expensive than a full binding calculation. In addition, emerging generative approaches such as BioEmu provide an alternative means of sampling approximate equilibrium protein ensembles and could be used to generate or characterise the folding component in future implementations (Lewis et al., 2025). Second, the method requires collective variables that adequately distinguish the relevant conformational and binding states. OneOPES combines enhanced-sampling strategies and can improve sampling even when the selected variables do not fully describe the slow dynamics. Recently suggested algorithm, developed for folding simulations and focusing of native and non-native contacts can automate CV selection (Rizzi et al., 2025). Yet, convergence must still be demonstrated explicitly (Rizzi et al., 2023). Third, our binding calculations employed funnel restraints and Cα-based restraints, whose effects needed to be accounted for explicitly in the free-energy calculation (Limongelli et al., 2013). Finally, we assumed that the receptor structure remained sufficiently stable during binding. For systems in which both binding partners undergo extensive conformational rearrangements, such as binding between two intrinsically disordered proteins, additional thermodynamic components may be required to account for receptor reorganization and co-folding.

Overall, our results show that coupled folding and binding in some cases need not be treated as a single high-dimensional sampling problem when calculating equilibrium free energies. By separating the folding penalty from the binding contribution, we reduced the conformational complexity of individual calculations while retaining the correct thermodynamic connection between the unbound and bound states. This strategy therefore provides a general framework for applying enhanced-sampling free-energy calculations to flexible peptides and disordered binding motifs, while allowing the underlying folding and binding mechanisms to be investigated independently.

### Conclusions

In this work, we demonstrated that novel enhanced-sampling approaches can yield structural, thermodynamic, and mechanistic insight into the coupled folding and binding of intrinsically disordered peptides. We showed that Circuit Topology provides a complementary framework for characterising conformational transitions in IDPs, while OneOPES enables efficient exploration of their conformational landscapes and the calculation of the absolute free energy. Our results further suggest that folding and binding can be treated as separate thermodynamic components, providing a practical strategy for free-energy calculations of highly flexible binding partners. At the same time, the coexistence of induced-fit and conformational-selection pathways highlights the complexity of the underlying binding process and the need for approaches that can resolve its kinetics. Future developments in enhanced sampling, kinetic modelling, and simulation-based prediction will be important for moving from thermodynamic descriptions to quantitative predictions of coupled folding-and-binding mechanisms.

## Data and Code Availability

The code developed and used in this study is available in the following GitHub repository: https://github.com/TheMashaghiLab/Publication_Multimodal_conformational_landscape_and_thermodynamics_of_-folding_and_binding_in_an_IDR

## Author contributions

A.M. and V.A. conceived the project and designed the research. V.A. performed the simulations and conducted the investigation. V.A., V.R., and A.M. developed the methodology. V.A. analysed the data. A.M. and J.v.N. acquired funding. A.M. provided computational resources, supervised the research, and administered the project. V.A., A.M., J.v.N., and V.R. reviewed the manuscript. V.A., A.M., and J.v.N. wrote the manuscript, with input from all authors.

