## Supplementary Information for "Multimodal conformational landscape and thermodynamics of coupled folding and binding in an intrinsically disordered protein region"

### Supplementary Methods:

#### Molecular dynamics

The structure of the TFIIF RAP74 and the RNAP II CTD Phosphatase FCP1 complex deposited in the Protein Data Bank (Berman et al., 2000) with code 1ONV was used as the initial structure for the simulation (Nguyen et al., 2003). We generated a structure for the isolated FCP1term simulation with 5 ns at 1000 K using the helical conformation of FCP1term from 1ONV. The final frame of the 1000 K MD was used as the initial structure for folding simulations.

The MD simulations were performed using the free molecular simulation software GROMACS version 2023.5 (Abraham et al., 2015) patched with the open-source, community-developed PLUMED (Bonomi et al., 2019) library version 2.9.3 (Tribello et al., 2014). The protein force field was DES-Amber (Piana et al., 2020) with explicit solvent; charges were scaled to 0.9, as suggested by force field developers. Proteins were solvated in a dodecahedron box with periodic boundary conditions. The box vector for the isolated $FCP1_{term}$ protein simulation was 6 nm × 6 nm × 6 nm. The box vector for the isolated Rap74:$FCP1_{term}$ complex simulation was 8 nm × 8 nm × 8 nm. The system was neutralized with 150 mM sodium and chloride ions. Before production runs, the system was minimised and equilibrated. The equilibration was performed in the NVT ensemble, followed by the NPT ensembles for 100 ps. The heavy protein atoms were restrained using a harmonic potential with the default constant value of 1000 kJ $mol^{-1}$ $nm^{-2}$. The LINCS algorithm (Hess et al., 1997) was used to constrain H-bond vibrations. We integrated using the leapfrog algorithm with a time step of 2 fs. Buffered neighbor searching was done with a Verlet list cutoff scheme, with a cutoff of 1.2 nm. Particle mesh Ewald (Darden et al., 1993) with cubic interpolation was used for long-range electrostatics. The pressure was maintained at 1 bar using a Parrinello–Rahman barostat, with a relaxation time of 2 ps for equilibration and 5 ps for production. The isothermal compressibility of water was $4.5 \times 10^{-5}$ $bar^{-1}$. We coupled the solute and solvent independently; we set the temperature to 300 K or 350 K, depending on the experiment. During equilibration, we used a modified Berendsen thermostat with a time constant tau of 0.1 ps. In production runs, we controlled the temperature with a Nose–Hoover thermostat, with a time constant tau of 1 ps.

All conventional simulations were done in a single run lasting 10 µs. Exploratory OPES_EXPLORE simulations were performed in four independent repeats, each lasting 8 µs. OneOPES simulations were performed for four independent repeats, each comprising eight replicas simulated for 500 ns.

**PLUMED analysis**

Contacts between residue pairs and atoms were calculated using the following formula:

$$S(r) = \frac{1-\left(\frac{r}{r_0}\right)^6}{1-\left(\frac{r}{r_0}\right)^8}, (1)$$

where r is the distance between the centers of mass of the side-chain heavy atoms of the residue pair for side-chain contacts, and r0 = 0.7 nm. We used the same formula to calculate helical contacts, where r is the distance between the backbone CO oxygen atom and the NH hydrogen that forms a hydrogen bond, and r0 is 0.3 nm. The list of interchain and intrachain contacts can be found in Supplementary Table 1.

To probe the helical content of $FCP1_{term}$ during MDs, we used the PLUMED internal CV ALPHARMSD (Pietrucci & Laio, 2009):

$$S_{helix} = \sum_i \frac{1-\left(\frac{r_i}{r_0}\right)^8}{1-\left(\frac{r_i}{r_0}\right)^{12}}, (2)$$

where $r_i$ is the RMSD distance from reference α-helical configuration, $r_0$ = 0.08 is the cut-off distance, summation goes over atoms of the selected region that are characteristic for helix formation.

**On-the-fly probability-enhanced sampling (OPES) method**

To investigate slow dynamics, we used On-the-fly probability-enhanced sampling (Invernizzi & Parrinello, 2020). During the simulation run, this method applies a bias along selected collective variables to achieve convergence in the reweighted ensemble. In this paper, we used a modification of this method, called OPES Explore (Invernizzi & Parrinello, 2022), designed to increase the exploration of the selected CVs space. In this modification, the aim is to achieve convergence of the target ensemble, which is chosen as a well-tempered ensemble:

$$p^{tg}(s) = p^{WT}(s) \propto [P(s)]^{1/\gamma} \text{ (3)},$$

where $p^{tg}(s)$ is a target distribution, $P(s)$ is the unbiased marginal distribution, s are the chosen CVs, and γ is the bias factor that controls the flattening of the distribution. Then the bias $V_n$ at each iteration *n* is calculated on the fly:

$$V_n = (\gamma - 1)k_B T \log\left(\frac{p_n^{WT}(s)}{Z_n} + \epsilon\right), (4)$$

where $k_B$ is the Boltzmann constant, *T* is the system temperature, $p^{WT}{}_n(s)$ is the well-tempered distribution at bias update step *n*, $Z_n$ normalises $p^{WT}{}_n(s)$ over the sampled CV space, and $\epsilon$ is a regularisation term that prevents infinite bias.

OPES has an additional variation called OPES Expanded. In this variation, the selected thermodynamic variable converges as the simulated system evolves. In our work, we used potential energy, scaled over a temperature range with MULTITHERMAL CV. Unlike conventional replica-exchange methods that require simulating many parallel replicas at different temperatures, OPES Expanded constructs a single adaptive bias potential along thermodynamic degrees of freedom, allowing a single simulation to sample a multicanonical target distribution across a defined temperature range.

In the exploratory run, we used OPES Explore with two CVs: $LDA_{interchain}$ – native contacts between two protein chains; and $LDA_{helix}$ – backbone hydrogen bonds forming an alpha helix in $FCP1_{term}$. We calculated these contact weights from two 300 ns conventional runs. In the first, we used the native complex structure (1ONV). In the second, $FCP1_{term}$ and Rap74 were detached. $FCP1_{term}$ was in the disordered state, as described above. During this simulation, we obtained statistics for the contact distribution (equation 1). Then, using the linear discriminant analysis module of scikit, we calculated the importance of each residue pair. In the exploratory runs, PACE was 1000, BARRIER 40 kJ/mol, and SIGMA for $LDA_{interchain}$ was 0.15, and for $LDA_{helix}$ was 0.1.

**On-the-fly Probability Enhanced Sampling with Replica Exchange (OneOPES) simulation protocol**

To achieve statistical convergence in the presence of suboptimal collective variables, enhanced sampling simulations were conducted using the On-the-fly Probability Enhanced Sampling with Replica Exchange (OneOPES) protocol (Rizzi et al., 2023). OneOPES combines multidimensional OPES Explore biases with OPES MultiThermal sampling across a stratified hierarchy of N = 8 replicas. All replicas share a common leading (primary) bias potential deposited on the primary CVs, which establishes a baseline for sampling. The first replica contains only this primary bias and serves as the convergence-focused replica from which we calculate equilibrium free energy surfaces (FES) via standard reweighting.

Higher-order replicas (replicas 2–8) incorporate an "exploration gradient" by progressively introducing weaker, slowly updated auxiliary biases on complementary CVs to accelerate transversal degrees of freedom. In addition, higher-order replicas use OPES MultiThermal to enable multithermal sampling and lower kinetic barriers. Coordinate exchanges between adjacent replicas are attempted at fixed intervals following the Metropolis–Hastings acceptance criterion.

**Conformational ensemble of isolated $FCP1_{term}$ in solution**

To reconstruct the free energy surface of unbound FCP1term in explicit solvent, OneOPES simulations were performed biasing the peptide helicity $S_{helix}$ as the main CV. We employed 8 replicas without additional supporting CV biases. Biased with BARRIER = 10 kJ/mol, PACE = 100000 timesteps, and a fixed kernel width of SIGMA = 0.015. We applied to Replicas 2–8 with upper temperature boundary array TEMP_MAX = [305, 310, 317, 327, 340, 355, 373]. Coordinate swaps were attempted every 10000 steps.

**Binding of helical $FCP1_{term}$ to RAP74**

To model the physical association of FCP1term pre-restrained in a helical conformation to the binding groove of RAP74, Funnel Metadynamics (Limongelli et al., 2013) was integrated into the OneOPES framework to restrict sampling to the relevant binding volume.

The simulation setup comprised three primary CVs: $S_N$ – fraction of native interchain contacts, $S_F$ – fraction of interchain contacts corresponding to the alternative/flipped pose and projection on axis **z** along the funnel. The residues used for defining $S_N$ and $S_F$ CVs can be found in Supplementary Table 1.

A funnel connecting the binding pocket to the solvent was created using the Cα atom of Val459 and the centre of mass of the Cα atom of Thr470 and the Cβ atom of Leu493, forming the z-axis. The funnel boundary was enforced using a rotationally symmetric potential with a sigmoidal profile:

$$V(s) = h/\left[1 + \exp\left(b(s - s_c)\right)\right] + f, \quad (5)$$

where h = 1.8 nm is the funnel radius at the binding pocket, f = 0.3 nm is the funnel radius in buffer, b = 2.5 $nm^{-1}$ is the slope, and sc = 3.0 is the inflection point along the funnel axis, **z**. To preserve domain integrity, an upper wall (UPPER_WALLS) restraint was applied to the Cα RMSD of RAP74 (a = 0.14 nm, *k* = 20000 kJ/mol). Backbone hydrogen-bond distance restraints were applied to $FCP1_{term}$ to maintain its helical structure throughout the trajectory (a = 12 nm, *k* = 400 kJ/mol) (Supplementary Table 1).

The main CVs were sampled with BARIER 50 kJ/mol, kernel widths SIGMA 0.02 ($S_N$), 0.02 ($S_F$),0.05 (projection to axis z) and PACE 100000. We also used a supporting CV, radial distance from axis z, with parameters: PACE 200000, BARIER 3 kJ/mol, SIGMA 0.02. Additionally, we added a bias region to exclude out-of-funnel restraints.

We used the same OPES MultiThermal component as in $FCP1_{term}$ in isolation OneOPES MDs.

**Coupled Folding and Binding of Unrestrained $FCP1_{term}$ to RAP74**

To simulate the fully unconstrained coupled folding and binding process, the backbone hydrogen-bond restraints on FCP1term were removed, allowing complete conformational freedom.

In the OneOPES simulations without restraints on the $FCP1_{term}$ helicity, we used the same main and supporting CVs and the same OPES MultiThermal component. However, to enhance peptide conformational transitions, we added $S_{helix}$ as the first supporting CV, starting from replica 2. Parameters for the $S_{helix}$ CV bias were: PACE 200000, BARRIER 5 kJ/mol, SIGMA 0.015. Starting from replica 3, we used the same parameters as above.

Additionally, we added restraints on $S_N$ and $S_F$, so that they could not decrease below 0.1 (a = 0.1, k = 10000 kJ/mol). This prevented over-biasing of the completely detached states.

We used exactly the same OPES MultiThermal component as in the OneOPES simulations of isolated FCP1term.

### Supplementary Results 1: Reweighted OneOPES ensemble of FCP1term replicates conventional MD ensemble

All four replicas showed a similar residue-resolved profile of helix propensity (Figure S2A). In contrast to the conventional simulation, the probability of observing residues in the helical state between residues 942–951 and 954–960 was similar, at around 10%. Taking into account the multiple replicas and the results of the block analysis of the conventional simulation, we estimate that the helical propensity of these regions is approximately 10%.

We then compared the FES along the helical collective variable coordinate between the OneOPES and conventional MD simulations (Figure S2B). The free-energy profiles from the OneOPES simulations and conventional MD are similar within approximately *k*BT (2.5 kJ/mol). All simulations show the same general behaviour: the completely disordered state is the most stable, followed by an initial rapid increase in free energy to 5–8 kJ/mol. At higher values of helical collective variable, the free energy continues to increase, reaching approximately 18 kJ/mol above the disordered state. One difference is that conventional MD shows a local minimum between $S_{helix}$ = 10 and 14, while the OneOPES simulations produce a much steeper free-energy surface in this region.

OneOPES repeat 3 showed a somewhat different free-energy profile. Starting from $S_{helix}$ = 6, its local minima are located approximately where the other simulations show maxima, and it reaches $\Delta F \approx 25$ kJ/mol at $S_{helix}$ = 13, compared with approximately 15 kJ/mol in the other simulations. Nevertheless, as discussed below, the residue-resolved distribution in this repeat remains consistent with the same general helix-formation mechanism.

Finally, to characterize the α-helix formation mechanism in the individual OneOPES repeats, we constructed the same 2D conditional probability histograms as for the conventional simulation (Figure S2C). Only repeat 3 showed a noticeably different mechanism. All other repeats showed residue-resolved probability distributions similar to those observed in the conventional simulation. At $S_{helix}$ values lower than 7, helices are preferentially formed around residues 942–951 and 954–960. The resulting histograms have a cone-like shape centred around Lys951. Incorporation of Lys951 into the helix is associated with formation of the extended helical structure. The histogram for repeat 3 is slightly shifted and asymmetric, but Lys951 remains close to the tip of the cone, suggesting that its role in the formation of the long helix is preserved.

It is important to note that the conventional MD and OneOPES simulation protocols showed good agreement, with the reweighted OneOPES ensemble producing a free-energy surface along the $S_{helix}$ CV that was highly similar to that obtained from conventional MD. Notably, comparable sampling of the $S_{helix}$ coordinate was achieved with approximately 4 μs of combined OneOPES simulations, compared with 10 μs of conventional MD. These results demonstrate that OneOPES can efficiently sample the conformational landscape of the disordered $FCP1_{term}$ peptide while reproducing the equilibrium ensemble obtained from substantially longer conventional MD simulations.

### Supplementary Results 2: NMR chemical shift analysis

Experimental NMR chemical shifts provide residue-specific information about conformational populations. To identify conformational populations contributing to the experimental NMR response, we calculated backbone NMR chemical shifts from the simulation ensembles using SPARTA+ (Shen & Bax, 2010) and compared them with previously reported experimental NMR data (Showalter & Showalter, 2009). We first calculated the NMR chemical shifts for the conformational ensemble generated by conventional MD (Figure S4). The predicted backbone chemical shifts indicated a highly disordered conformational ensemble for $FCP1_{term}$ in solution. After correcting for the random-coil contribution, the average calculated chemical shifts were close to zero (Figure S4A) and generally lower than the corresponding experimental values. However, the experimental chemical shifts largely fell within the standard deviation of the calculated values, suggesting that some conformations sampled during the simulations resemble structures present in the experimental ensemble.

To determine whether these results could arise from insufficient sampling in conventional MD simulations, we performed the same analysis using OneOPES simulations. NMR chemical shifts were calculated for both the reweighted ensemble (Figure S5) and the well-tempered ensemble (Figure S6). In the first case, we corrected for the bias when calculating averages; in the second, we did not. Interestingly, the reweighted ensemble produced chemical shift profiles similar to those obtained from conventional MD (Figure S5A). Across all repeats, the calculated chemical shifts remained close to zero after random-coil correction. Although the RMSE values were below 1 ppm, the Pearson correlation coefficients remained low, reaching a maximum value of 0.303 (Figure S5B). Thus, we concluded that the results were not dependent on the sampling method and that the same pattern was observed.

In contrast, a substantially different result was obtained for the well-tempered ensemble. A strong agreement between the calculated and experimental chemical shifts was observed across all repeats (Figure S6A). In all repeats, the RMSE did not exceed 0.413 ppm, while the lowest Pearson correlation coefficient was 0.738 (Figure S6B). These results indicate that the conformational populations sampled in the well-tempered ensemble agree considerably better with the experimental NMR data. The free-energy profiles further show that the barrier separating disordered and helical conformations is substantially lower in the well-tempered ensemble than in conventional MD (Figure S6C).

### Supplementary Results 3: Rap74 dynamics

Rap74 is well-folded in both the *apo* and *holo* states, adopting three stable α-helices (H1–H3) and one antiparallel β-sheet (Figure 3A). Helices H2 and H3 are connected by a flexible loop region, L. The

Rap74:FCP1term complex is stabilised by a network of hydrophobic (Figure 3B) and charged (Figure 3C) interactions. These interactions have been described in detail previously and are therefore not discussed further here (Kamada et al., 2003; Nguyen et al., 2003; Wostenberg et al., 2011).

Despite its overall structural stability, Rap74 in complex with $FCP1_{term}$ exhibited conformational fluctuations during the conventional MD simulations. The largest changes involved the loop connecting H2 and H3, which transiently adopted a helical conformation. At elevated temperature, we also observed additional changes in the regions surrounding H2 and H3, including a partial loss of helical structure. To characterize these conformational changes, we performed tICA using a lag time of 50 ps. We selected intrachain contacts with a variance greater than 0.02, thereby excluding contacts that remained essentially unchanged throughout the simulations. The resulting contact fluctuations were concentrated primarily in the loop region and the N-terminal disordered region of Rap74 (Figure S7A,B). Projection of the trajectories onto the first two tICA components revealed four distinct conformational clusters (Figure S7C).

We next examined the temperature dependence of the conformational dynamics by projecting the trajectories onto the first tICA component. The second tICA component did not provide additional separation of the identified states, as the four clusters could already be distinguished along tICA1; therefore, the subsequent analysis was performed using this coordinate alone (Figure S7D). Simulations at both 300 and 350 K were initiated from state 2. At 300 K, the system remained in state 2 for approximately 6 µs before transitioning to state 1, with no further transitions observed within the simulation time. At 350 K, transitions occurred more frequently, and the system repeatedly interconverted between the identified states.

The structural characteristics of these states explain the temperature-dependent behaviour (Figure S7E). At 300 K, states 1 and 2 were predominantly populated. In state 1, the loop between H2 and H3 adopted a helical conformation. At 350 K, increased conformational fluctuations resulted in partial loss of helical structure in the region surrounding H2, corresponding to state 3, and partial melting of H3, corresponding to state 4. Thus, increasing the temperature primarily affects the local structural elements of Rap74 while the overall fold remains preserved.

### Supplementary Results 4: Mechanism of folding-and-binding

#### FCP1term with no restrictions

OneOPES simulations without helical restraints provide insight into the mechanism of Rap74–FCP1term complex formation. The free-energy surfaces for repeats 2–4 (Figure S13A) are consistent with an induced-fit mechanism, in which FCP1term binds Rap74 before adopting its fully folded helical conformation. The free-energy gradients indicate that complete folding is not required to reach the native bound state. Instead, the peptide can initially form a subset of native contacts in a partially folded conformation, followed by the progressive formation of additional contacts and helical structure. In particular, no distinct low-free-energy basin is observed at high $S_{helix}$ (>10) and low $S_N$ (<0.1), as would be expected if a fully folded peptide were required for binding. Instead, an intermediate basin is observed at $S_{helix} < 6$ and $S_N < 0.8$. From this region, both native contacts and helical structure progressively develop towards the native state. The extension of the native-state basin along the $S_{helix}$ coordinate further indicates that binding and folding are coupled rather than strictly sequential processes. Importantly, these observations do not exclude an induced conformational selection mechanism.

However, FCP1term can also form a disordered complex with Rap74. In all repeats, we observed a local minimum corresponding to fully disordered peptide conformations ($S_{alpha} < 1$) with partial native binding ($0.2 < S_N < 0.8$; Figure S13A). These conformations represent a range of disordered but variably bound states, stabilised primarily by hydrophobic contacts with Rap74. Notably, these disordered bound conformations may also serve as intermediates along the folding pathway, as observed in replicate 3.

To determine which contacts form during the initial stages of binding, we performed a conditional probability analysis of the native contacts included in the $S_N$ CV (Figure S13B). For each binned value of $S_N$, we calculated the probability of observing each contact, taking the OneOPES bias into account. This analysis reveals the order in which individual native contacts become established. Across all four repeats, contacts involving Leu953 and Leu957 form at the earliest stage of native-contact formation ($S_N$ = 0.4–0.5). Contacts involving Met949 form subsequently, while most of the remaining native contacts become stably populated at $S_N > 0.7$. Contacts involving Leu954 and Asn958 behave differently. Although these contacts can form at $S_N > 0.3$, their conditional probabilities remain around 0.4, indicating that they are relatively dynamic and not consistently maintained during the early stages of binding.

We next examined $FCP1_{term}$ helix formation on the Rap74 surface. Complete helix formation occurs only at high $S_N$ values (>0.8), indicating that the fully folded helix forms late in the binding process. However, partial helical structure appears earlier. In repeats 2–4, residues 948–956 show a helix-formation probability above 0.4 at $0.7 < S_N < 0.8$, defining an early, relatively stable helical region. At lower values of $S_N$ (0.5–0.7), residues 952–955 are the first region to preferentially adopt a helical conformation. Thus, native-contact formation and helix formation overlap during binding, with the central part of the helix forming before the complete helical structure is established.

Figure 6 illustrates the proposed mechanism. Initially, Leu953 and Leu957 form hydrophobic contacts with Val490, Leu474, Leu493, and Leu497 (Figure 6A). At this stage, Leu957 has not yet penetrated deeply enough into the binding pocket to form all its native contacts. These initial interactions facilitate formation of a helical segment involving residues 952–957 (Figure 6B). Subsequently, Met949 forms additional interactions with Ser486 and Thr489, completing interactions with the corresponding pocket (Figure 6C). Finally, Leu957 moves deeper into the binding pocket and contacts Phe513, completing the native interaction network (Figure 6D). The complete structural transition is illustrated in Supplementary Movie 1. Because the trajectory shown in this movie was generated by demultiplexing OneOPES repeat 3, its temporal evolution should not be interpreted as representative of the kinetics of the binding process.

Overall, the simulations indicate that binding is initiated by the formation of hydrophobic contacts, accompanied by the formation of a short helical segment, followed by progressive helix stabilisation and completion of the native contact network. The initial contact formation and partial helix formation may also occur simultaneously when the peptide already samples a helical conformation before binding. An example of this alternative pathway is shown in Supplementary Movie 2.

#### Mechanism for the FCP1term with helical structure restriction

When the helical conformation of $FCP1_{term}$ is restrained, the binding process is consistent with a conformational-selection mechanism. The reweighted free-energy surfaces from the OneOPES simulations (Figure S14A) support this interpretation. At low $S_N$ (< 0.1) and high $S_{alpha}$ (> 10), the system occupies a high-free-energy region. A local minimum around $S_N \approx 0.3$ corresponds to the F conformation, whereas at S_N > 0.7, the free-energy surface contains minima corresponding to the N bound state.

Despite the helical restraint, the order of native-contact formation is like that observed for the unrestricted peptide. Contacts involving Leu953 and Leu957 form first, followed by those involving Met949, with the remaining native contacts established at higher $S_N$ values (Figure S14B). Thus, the initial stages of binding appear to involve a similar sequence of contact formation, regardless of whether the peptide is allowed to fold freely.

The simulations also revealed a substantial reorientation of the $FCP1_{term}$ helix within the binding pocket, including several complete transitions between the N and F orientations without peptide dissociation (Supplementary Movie 2). To characterise this motion, we used an orientation CV, defined earlier (Figure 3D, Figure S8A).

The folded helix progresses toward the native binding state more directly than the peptide undergoing folding on the Rap74 surface. Although we did not perform explicit kinetic modelling in this study, the FES shapes provide qualitative insight into the preferred transition directions. In Figure 6E, we compare the FESs along the binding coordinate for the helix-restrained and unrestricted $FCP1_{term}$ simulations. To focus on the native-binding pathway, we excluded configurations with SF > 0.4, which predominantly correspond to the flipped state. The resulting FES for the helix-restrained peptide decreases progressively toward the native state. In contrast, for the unrestricted $FCP1_{term}$ simulations, the average FES remains approximately flat for $S_N < 0.6$ and decreases for $S_N > 0.6$ as the peptide approaches the native state. These differences suggest that a pre-folded helical peptide has a greater thermodynamic tendency to progress toward the native binding state once it forms initial native contacts. In the unrestricted helix, the peptide can form initial contacts while remaining partially folded, resulting in a broader region of similar free energy before the final native contacts are established. Together, these FES profiles are consistent with the two limiting cases of coupled folding and binding: binding of a pre-formed helix followed by progression towards the native state, and binding accompanied by subsequent folding and the consolidation of native contacts.

The two-dimensional FESs projected onto $S_N$ and the orientation CV indicate that the helical peptide undergoes substantial rotational fluctuations within the binding pocket. In conventional MD simulations at 300 K, unbinding is associated with changes in helical orientation (Figure 3D), and this relationship is even more pronounced in the OneOPES simulations (Figure S14C). Following the free-energy gradients suggests that peptide unbinding is accompanied by rotation within the binding pocket. Importantly, these rotational motions can also drive transitions between the N- and F-bound states without requiring complete dissociation of FCP1term. Thus, the binding pocket allows the peptide to reorient between alternative bound configurations while maintaining the overall association with Rap74.

**Supplementary Table 1**

| CV name | Description | $r_0$, nm | Contact pairs |
|---|---|---|---|
| $S_N$, $LDA_{interchain}$ | Native state interchain contacts. The distance is calculated between centers of mass of the heavy atoms of the side chains. Numbers represent residue indices. First is the index of Rap74 aminoacid, second is the index of $FCP1_{term}$ aminoacid. | 0.7 | 493-953, 474-953, 490-953, 513-957, 497-957, 470-953, 502-957, 478-949, 489-949, 486-949, 501-960, 498-956, 471-947, 471-954, 504-958 |
| $S_F$ | Flipped state interchain contacts. The distance is calculated between centers of mass of the heavy atoms of the side chains. Numbers represent residue indices. The first number indicates the residue index in Rap74, and the second indicates the residue index in FCP1term. | 0.7 | 474-960, 490-961, 490-960, 490-957, 513-949, 497-953, 513-953, 498-954, 510-959, 504-948, 504-945, 498-947 |
| $LDA_{helix}$ | Native helical contacts between the backbone CO oxygen atom and NH hydrogen of the $FCP1_{term}$. The numbers represent atom serial numbers in the pdb. | 0.3 | 1188-1227, 1203-1242, 1213-1259, 1225-1269, 1240-1291, 1257-1301, 1267-1320, 1289-1335, 1299-1345, 1318-1360, 1177-1215, 1333-1379, 1343-1393, 1358-1405 |
| Helix restraints | Native helical contacts between the backbone CO oxygen atom and NH hydrogen of the $FCP1_{term}$. The numbers represent the atom serial numbers in the pdb. | 0.3 | 1188-1227, 1203-1242, 1213-1259, 1225-1269, 1240-1291, 1257-1301, 1267-1320, 1289-1335, 1299-1345, 1318-1360, 1177-1215, 1333-1379, 1358-1393 |

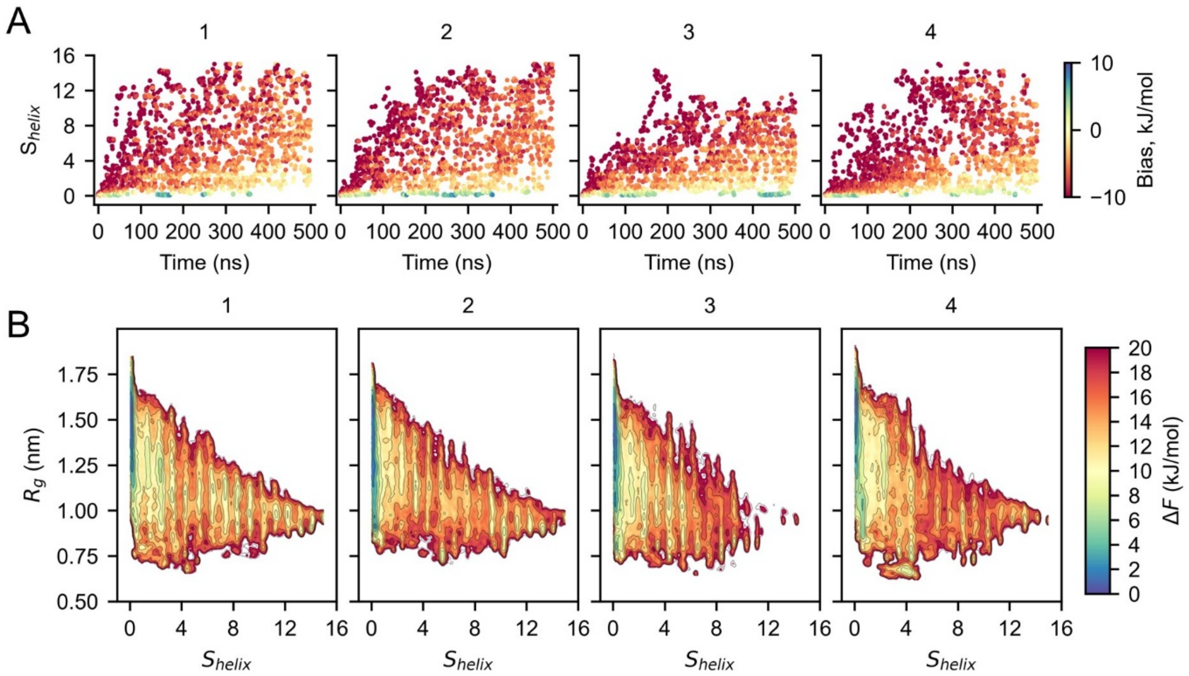


*Figure S1. OneOPES efficiently samples the conformational space of FCP1term. (**A**) Number of helical segments $S_{helix}$ as a function of simulation time for each independent repeat. Data are shown only for replicate 0. (**B**) Final reweighted FESs projected onto the $S_{helix}$ and $R_g$ coordinates for each independent repeat.*

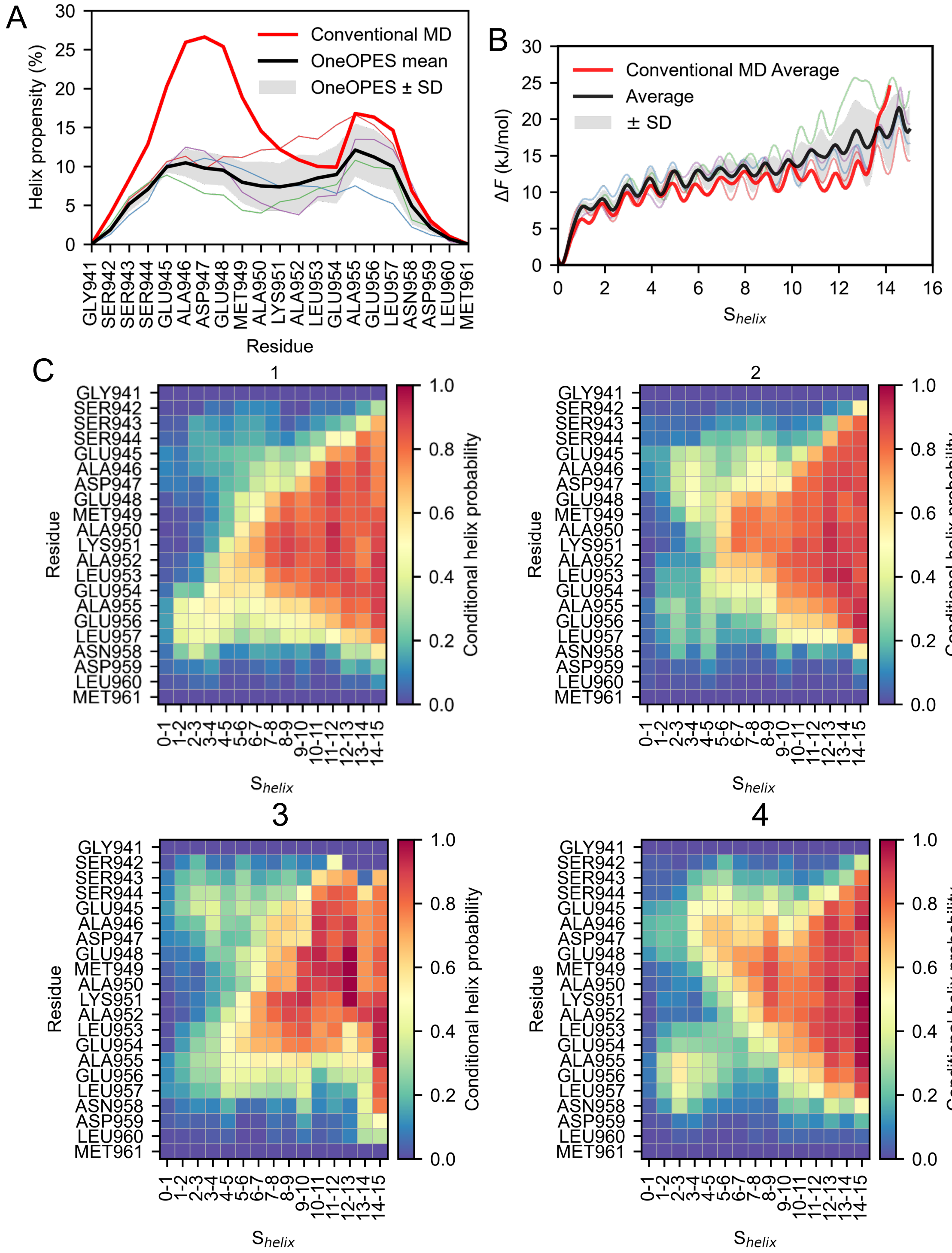


*Figure S2. Folding mechanism and the role of Lys951. (**A**) Helix propensity obtained from conventional MD (bold red line) and four independent OneOPES repeats (thin colored lines) from the reweighted ensemble. The average OneOPES helix propensity is shown as a bold black line, and the grey shaded area represents the standard deviation between repeats. (**B**) FES projected onto the $S_{helix}$ CV. The notation is the same as in (**A**). (**C**) Two-dimensional conditional helix-probability maps for each independent OneOPES repeat, calculated from the reweighted ensemble. The x-axis shows ranges of the $S_{helix}$ CV, while the y-axis shows residue numbers and amino acid identities. The color indicates the probability that a given residue adopts a helical conformation within the corresponding range of $S_{helix}$.*

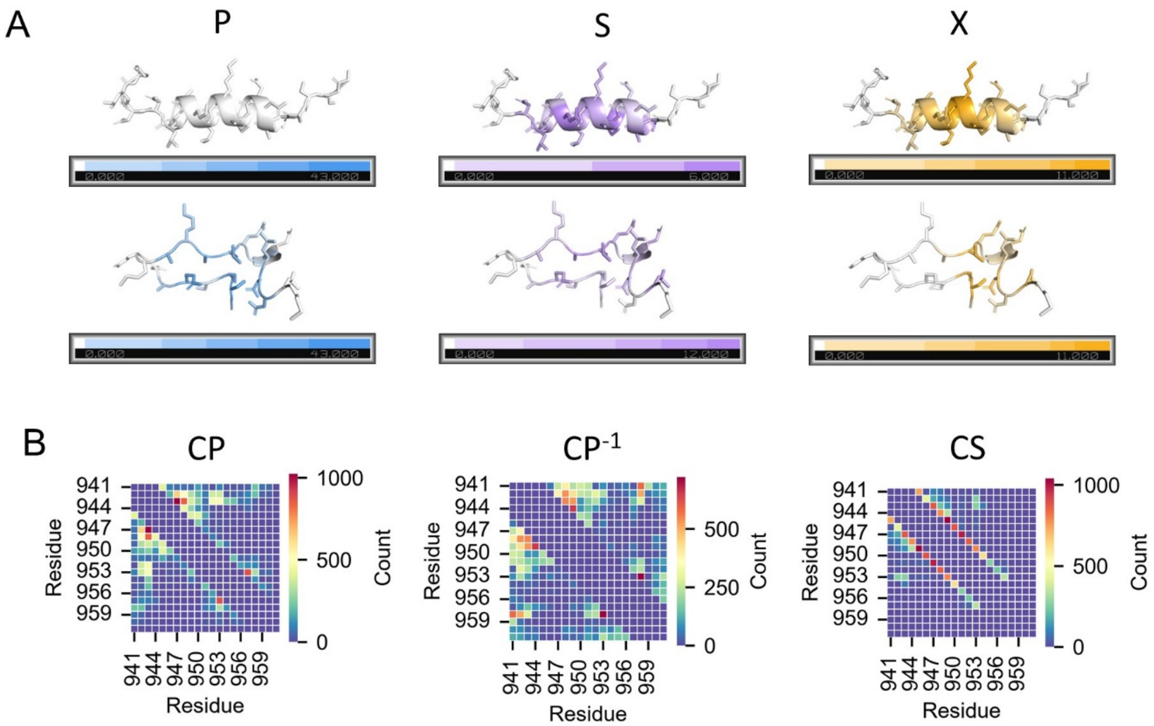


*Figure S3. Topological characteristics of the FCP1term peptide folding. (**A**) Distribution of parallel (P), series (S) and cross (X) contacts in the representative helix and beta sheet structures. (**B**) Average number of CP, $CP^{-1}$ and CS contacts per residue pair.*

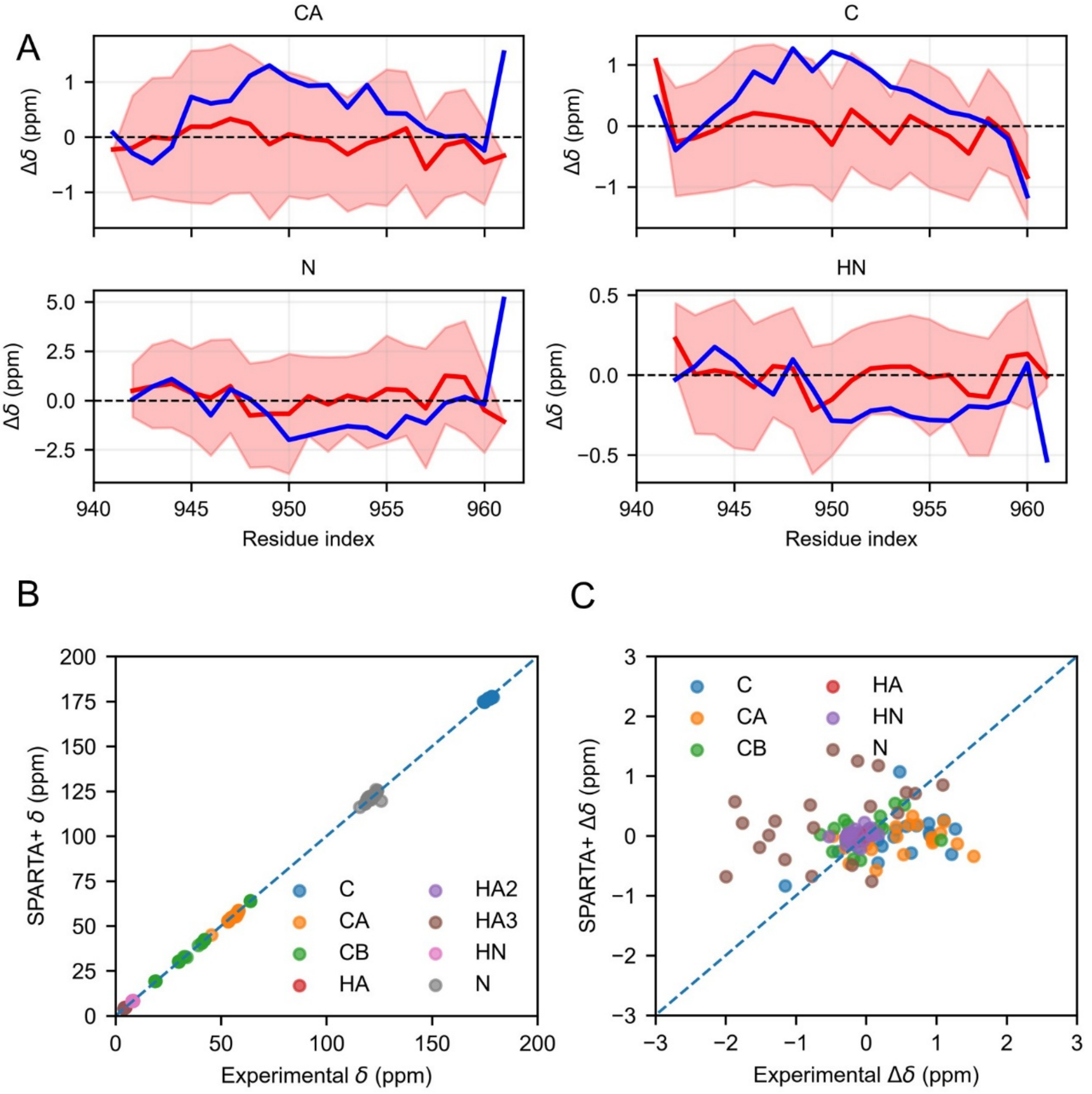


*Figure S4. Comparison of experimental and SPARTA+-calculated NMR chemical shifts for FCP1term. (**A**) NMR chemical shifts are plotted as a function of residue index for protein backbone atoms: Cα (CA), carbonyl carbon (C), backbone nitrogen (N), and amide proton (NH). All chemical shifts were corrected by subtracting the corresponding random-coil chemical shifts. The red line represents the average chemical shifts calculated from the conformational ensemble generated using conventional MD, while the red shaded area indicates the standard deviation of the calculated shifts. Experimental chemical shifts are shown as a blue line. (**B**) Comparison between calculated and experimental NMR chemical shifts without correction for the random-coil contribution. (**C**) Comparison between calculated and experimental NMR chemical shifts after correction for the random-coil contribution. The agreement between calculated and experimental shifts is characterized by an RMSE of 0.708 ppm, and a Pearson correlation coefficient of 0.193.*

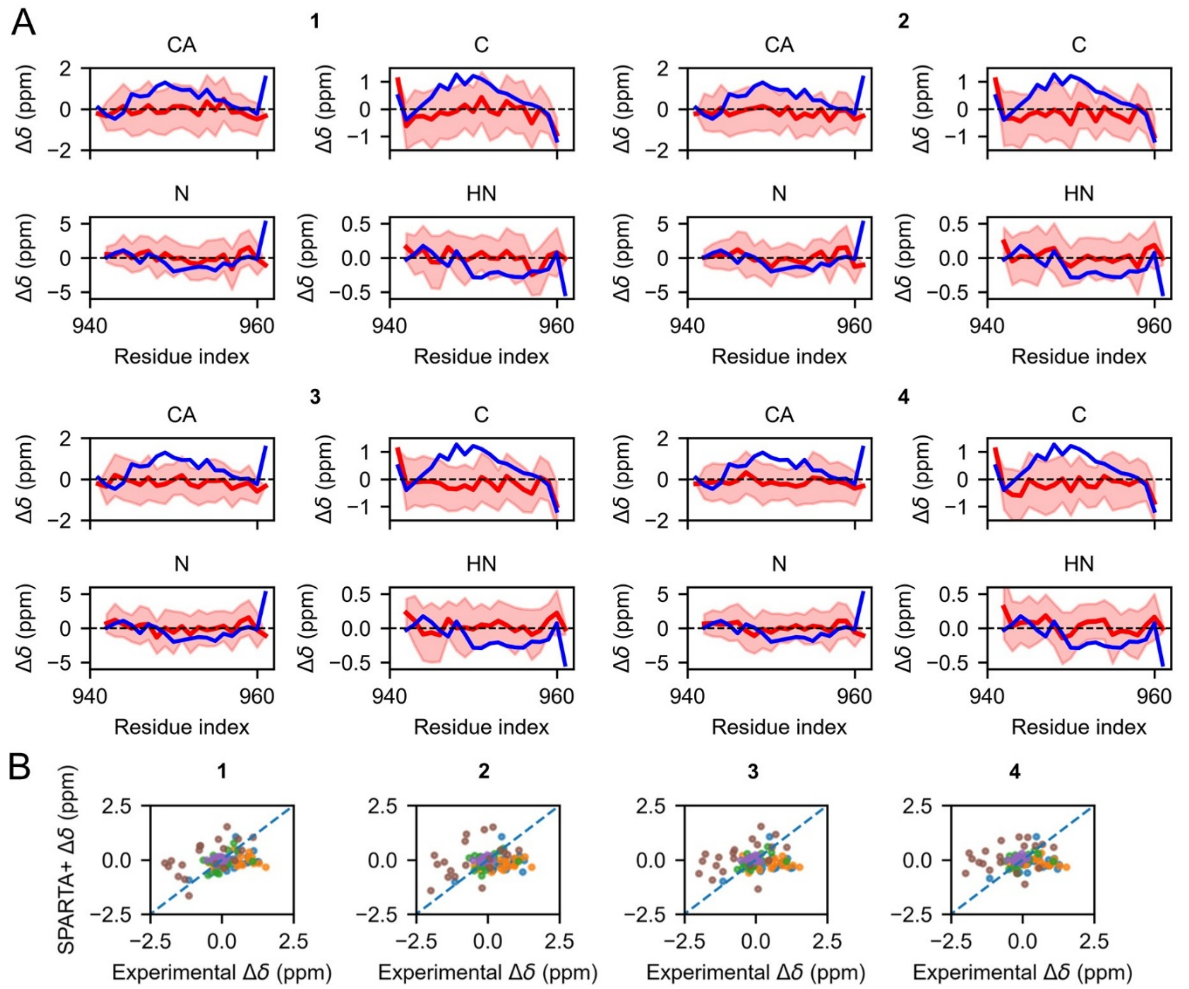


*Figure S5. Comparison of experimental and SPARTA+-calculated NMR chemical shifts for FCP1$_{term}$ from OneOPES simulations. (A) NMR chemical shifts, same as in Figure S4A, plotted for each repeat. The red line represents the average chemical shifts calculated from the reweighted conformational ensemble generated using OneOPES protocol, while the red shaded area indicates the standard deviation of the calculated shifts. Experimental chemical shifts are shown as a blue line. (B) Comparison between calculated and experimental NMR chemical shifts after correction for the random-coil contribution for each repeat. Agreement between calculated and experimental chemical shifts for each repeat: 1 – RMSE 0.638 ppm, Pearson correlation coefficient 0.303, 2 – RMSE 0.686 ppm, Pearson correlation coefficient 0.187, 3 – RMSE 0.691 ppm, Pearson correlation coefficient 0.130, 4 – RMSE 0.706 ppm, Pearson correlation coefficient 0.033.*

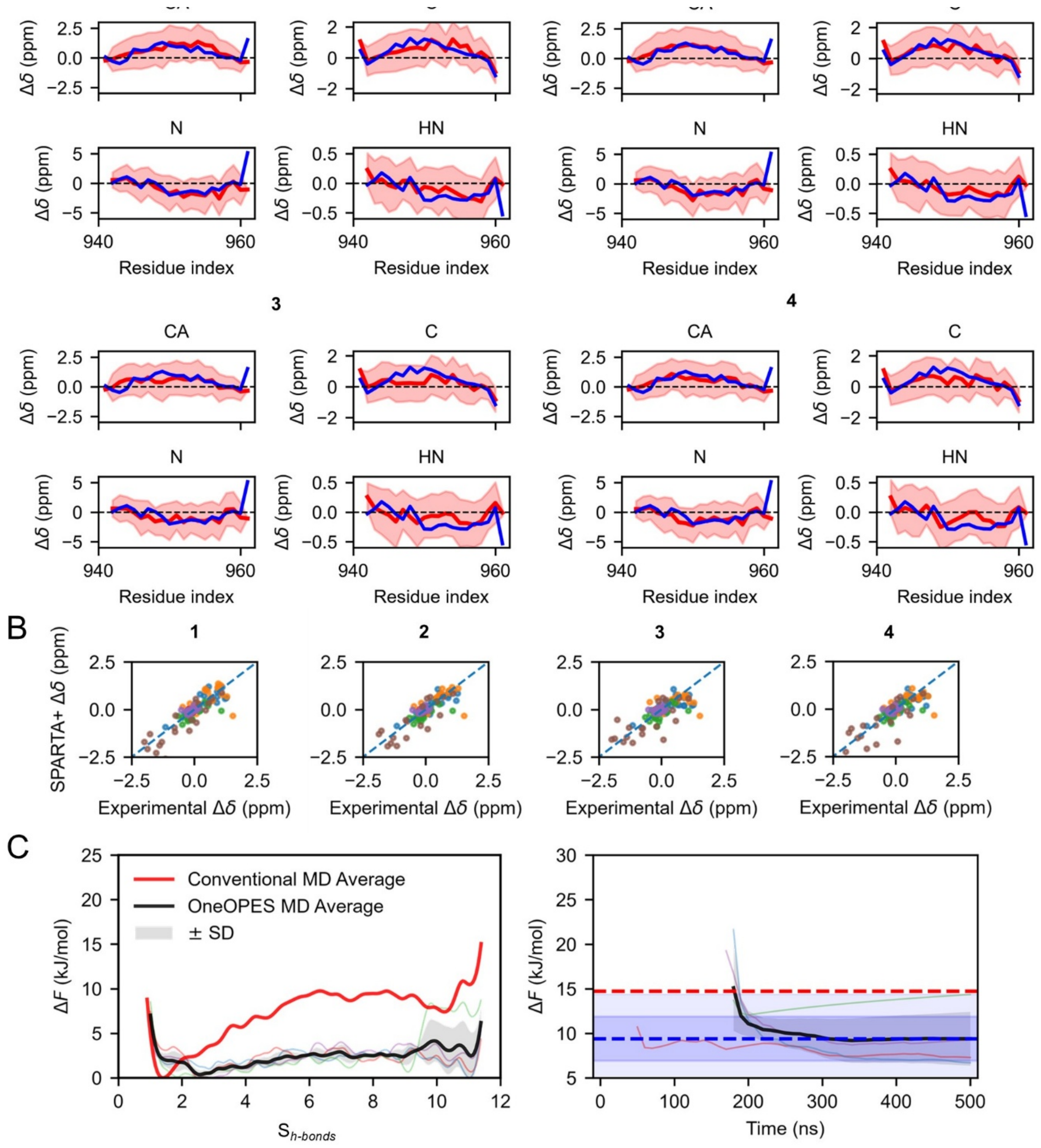


*Figure S6. Comparison of experimental and SPARTA+-calculated NMR chemical shifts (A) NMR chemical shifts, as described in Figure S4, are plotted for each repeat. In contrast to the analysis shown in Figure S4, the calculated chemical shifts were obtained without applying bias reweighting. Therefore, the calculated NMR chemical shifts represent the converged well-tempered ensemble. (B) Comparison between calculated and experimental NMR chemical shifts, corrected for the random-coil contribution, for each repeat. The agreement between calculated and experimental chemical shifts was evaluated separately for each repeat: repeat 1, RMSE = 0.388 ppm and Pearson correlation coefficient = 0.837; repeat 2, RMSE = 0.359 ppm and Pearson correlation coefficient = 0.850; repeat 3, RMSE = 0.443 ppm and Pearson correlation coefficient = 0.738; repeat 4, RMSE = 0.413 ppm and Pearson correlation coefficient = 0.782. (C) Left - Free-energy surfaces projected along the $S_{h\text{-}bonds}$ CV for conventional MD (bold red) and individual OneOPES repeats in the well-tempered ensemble (coloured thin lines). The average free-energy surface across all repeats is shown in black, and the standard deviation between*

*repeats is indicated by the grey shading. Right – convergence of the free energy difference. Notation same as in Figure 5F. Final OneOPES* $\Delta F_{folding}$ *= 9 ± 3 kJ/mol.*

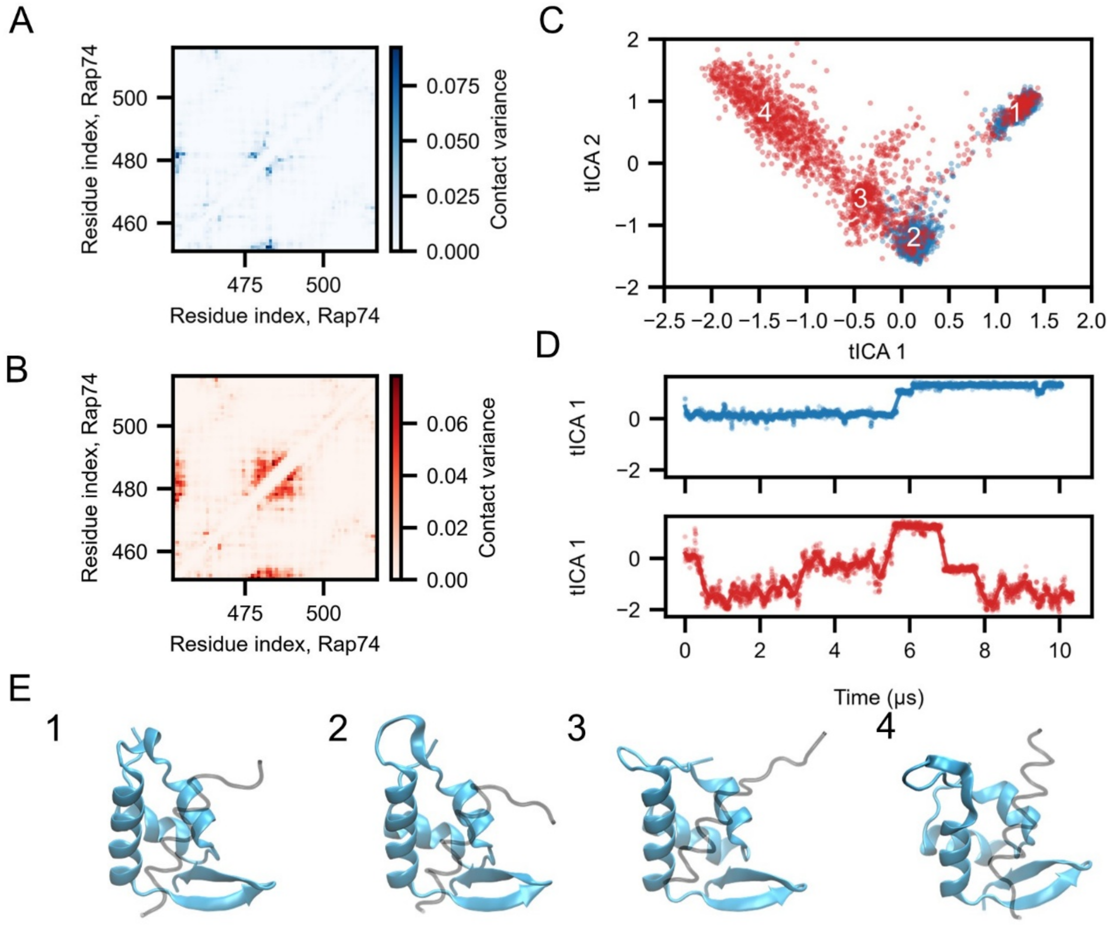


*Figure S7. Dynamics of the Rap74 structure. Variance of intrachain Rap74 contacts during conventional MD simulations at 300 K (**A**) and 350 K (**B**). (**C**) Conformational distribution of the Rap74:FCP1term complex projected onto the first two tICA axes. Four distinct conformational states are identified. (**D**) Time evolution of the tICA collective variables during the simulations. (**E**) Representative structures of the identified conformational clusters.*

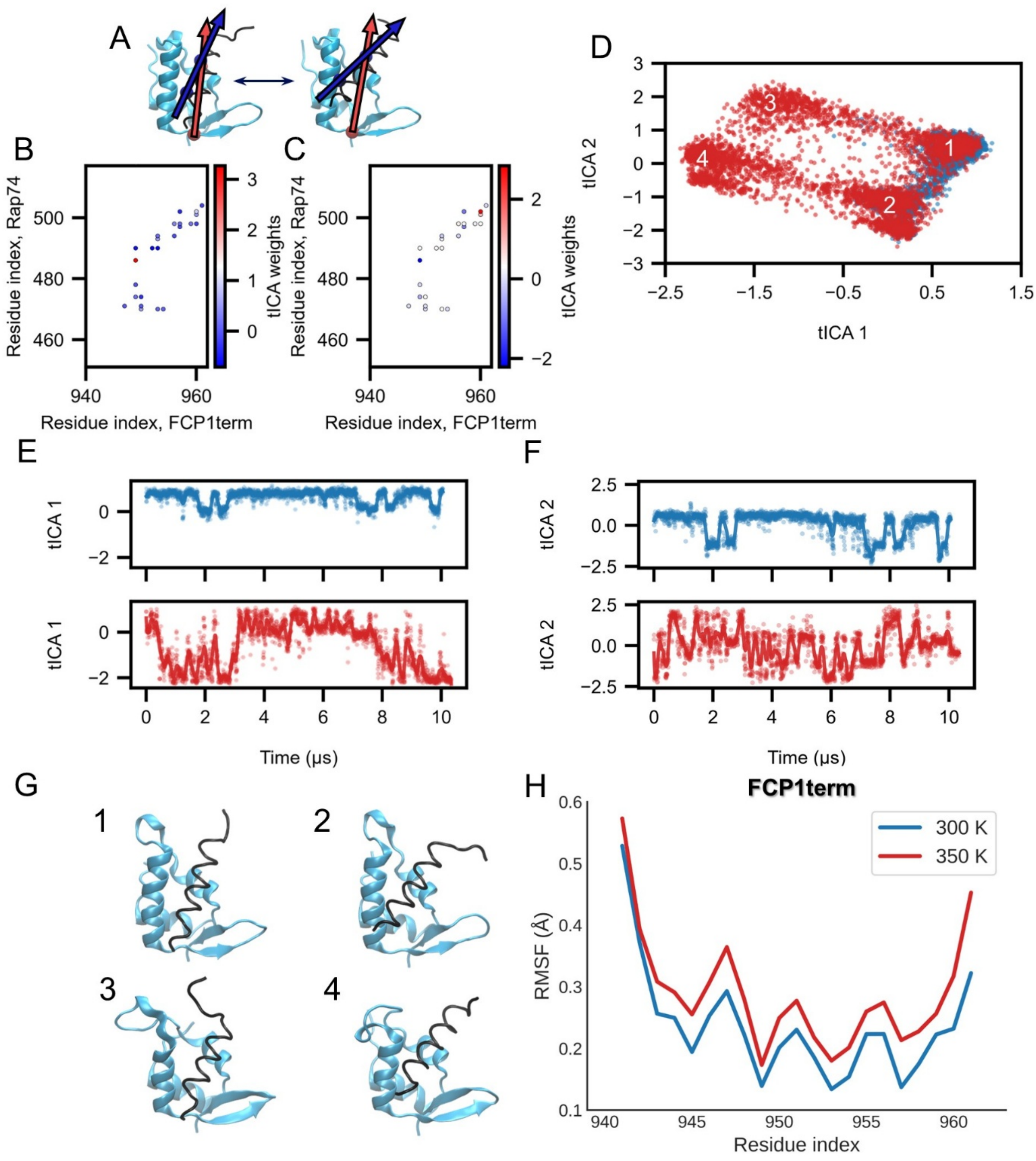


*Figure S8. FCP1term dynamics in the binding pocket. (**A**) Vectors defining the orientation of FCP1term (red) in the binding pocket of Rap74 (blue). The pocket vector is defined from the Cα atom of Pro502 to the C atom of Leu474, while the helix vector is defined from the Cα atom of Glu956 to the N atom of Met949. (**B**,**C**) Interchain contact wights for the first and second tICA components, respectively. (**D**) Conformational distribution of the Rap74:FCP1term complex projected onto the tICA axes. (**E**, **F**) Time evolution of the tICA collective variables at 300 K (blue), and 350 K (red). (**G**) Representative structures of the identified conformational clusters. (**H**) Root mean square fluctuation RMSF of the Cα atoms of FCP1term at 300 K (blue), and 350 K (red).*

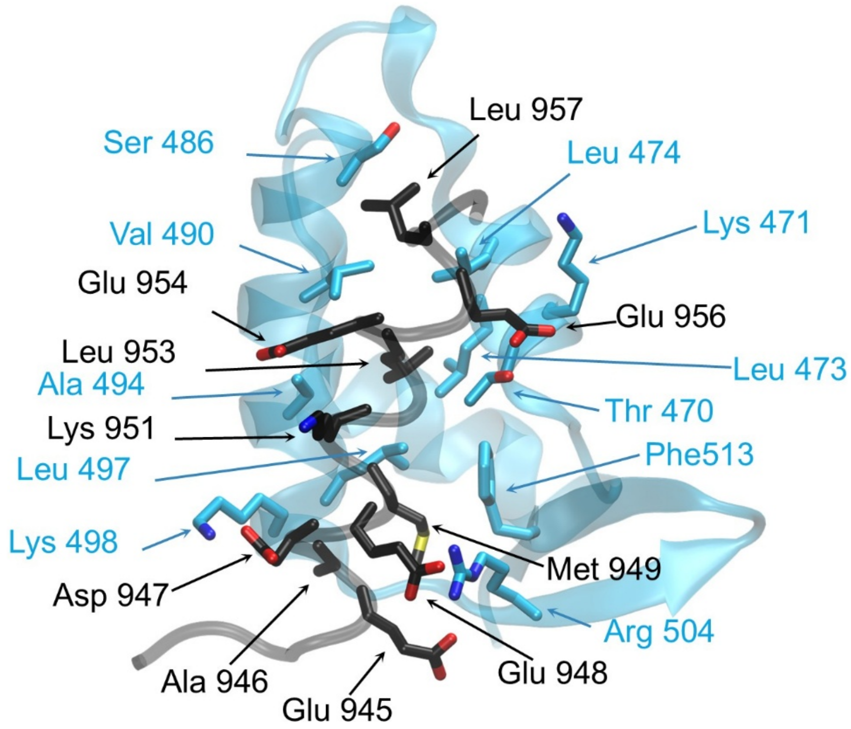


*Figure S9. Structure and contacts formed between FCP1term (black) and Rap74 (cyan)*

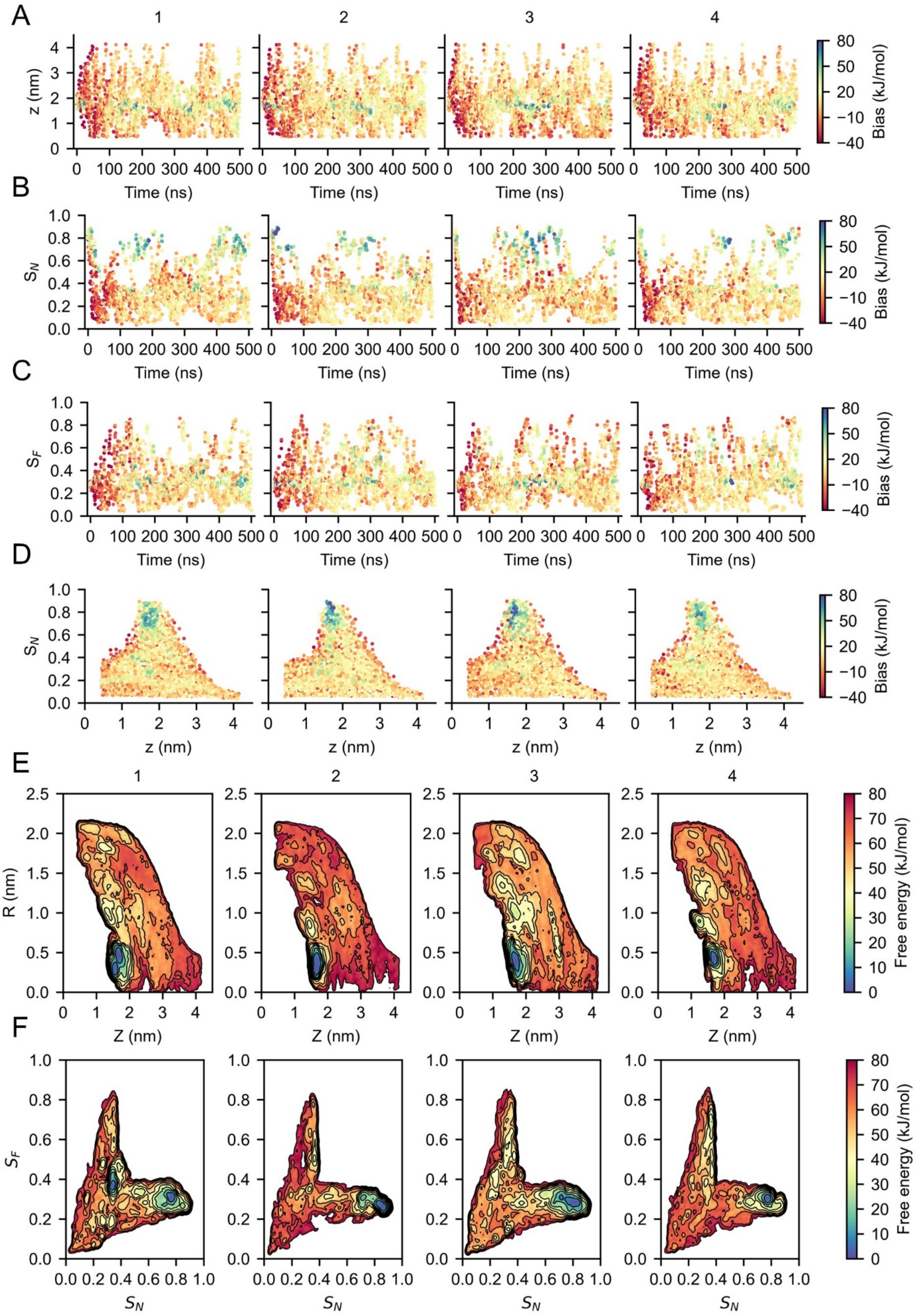


*Figure S10. Sampling of OneOPES simulations of Rap74:FCP1$_{term}$ system with helix restriction. Dynamics of center of mass distance (**A**), native (**B**) and flipped (**C**) contacts in the first replica for all repeats, color*

*represents the bias. (**D**) Distribution of distance from binding pocket and native contacts per in the first replica per repeat, color represents the amount of bias. (**E**) Reweighted FES projected to radial distance R and distance from the pocket for each independent OneOPES repeat. (**F**) Reweighted FES projected to $S_N$ and $S_F$ axis for each independent OneOPES repeat.*

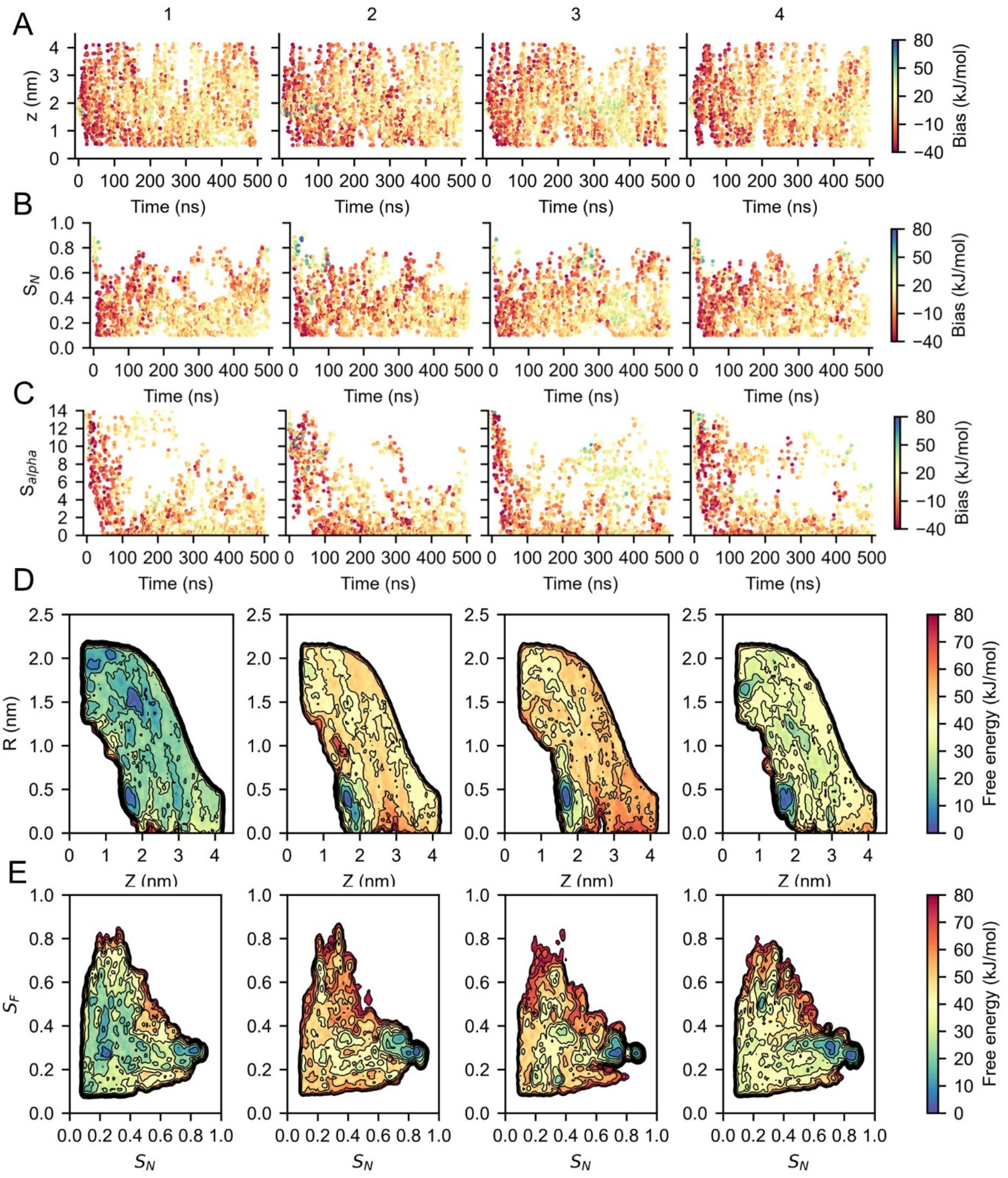


*Figure S11. Sampling of OneOPES simulations of Rap74:FCP1$_{term}$ system. (**A-C**) Time series of z, $S_N$, and peptide helicity $S_{alpha}$, respectively, for the first replica of each independent repeat. Color indicates the applied bias. (**D**) Reweighted FES projected to radial distance R and distance from the pocket for each independent OneOPES repeat. (**E**) Reweighted FES projected to $S_N$ and $S_F$ axis for each independent OneOPES repeat.*

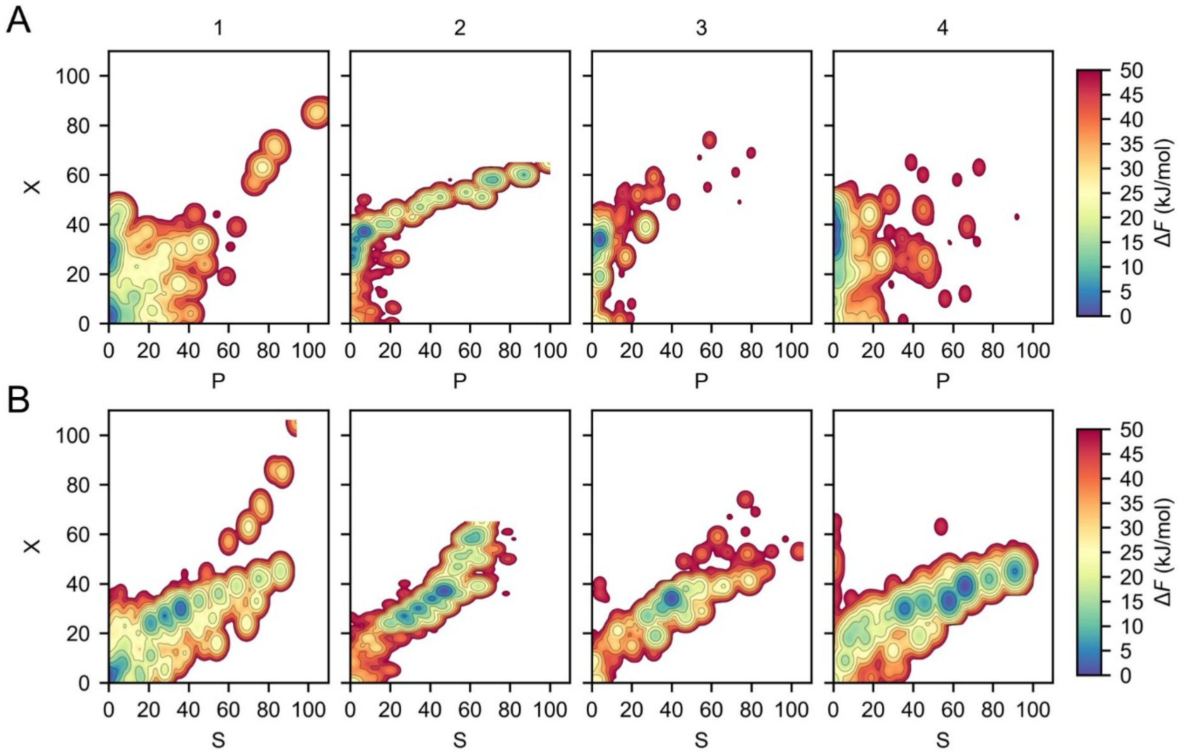


*Figure S12. FCP1$_{term}$ circuit topology organization in the presence of Rap74. (**A**) Reweighted free-energy surfaces projected onto P and X coordinates for each independent OneOPES repeats. (**B**) Corresponding surfaces projected onto S and X coordinates. In repeats 2–4, the free-energy minimum lies at X>20 and P<20, what represent a helical organization of the peptide.*

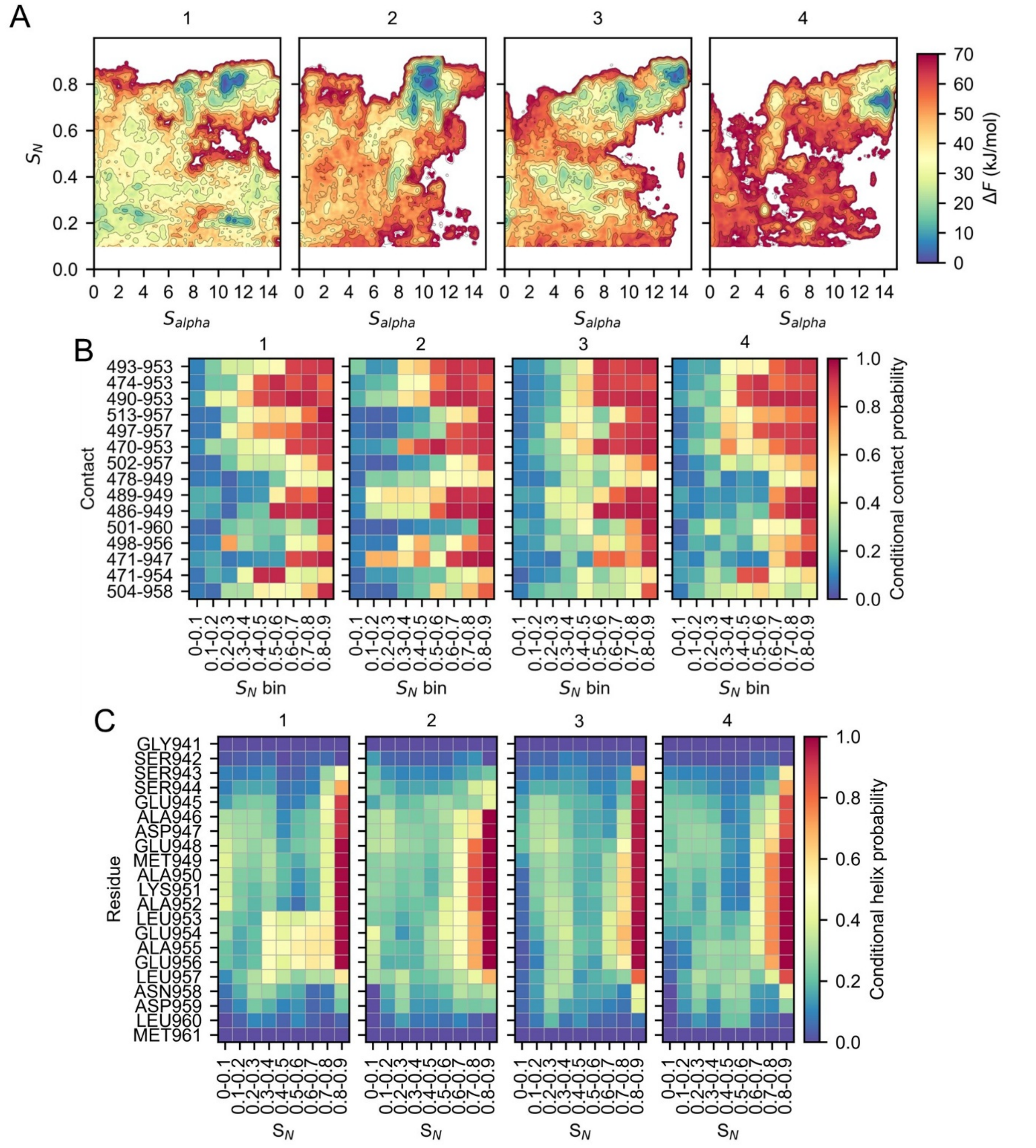


*Figure S13. Induced fit mechanism. (**A**) Reweighted free-energy surfaces projected onto the $S_{alpha}$ and $S_N$ coordinates for each independent OneOPES repeat. (**B**) Two-dimensional conditional contact-probability maps for each independent OneOPES repeat, calculated from the reweighted ensemble. The x-axis shows ranges of the $S_N$ CV, while the y-axis lists the residue pairs forming interchain contacts. For each pair, the first index corresponds to the Rap74 residue and the second to the FCP1$_{term}$ residue. The color indicates the probability of observing the corresponding contact within each $S_N$ range. (**C**) Conditional probability that each residue adopts a helical conformation, binned by $S_N$, for each independent repeat.*

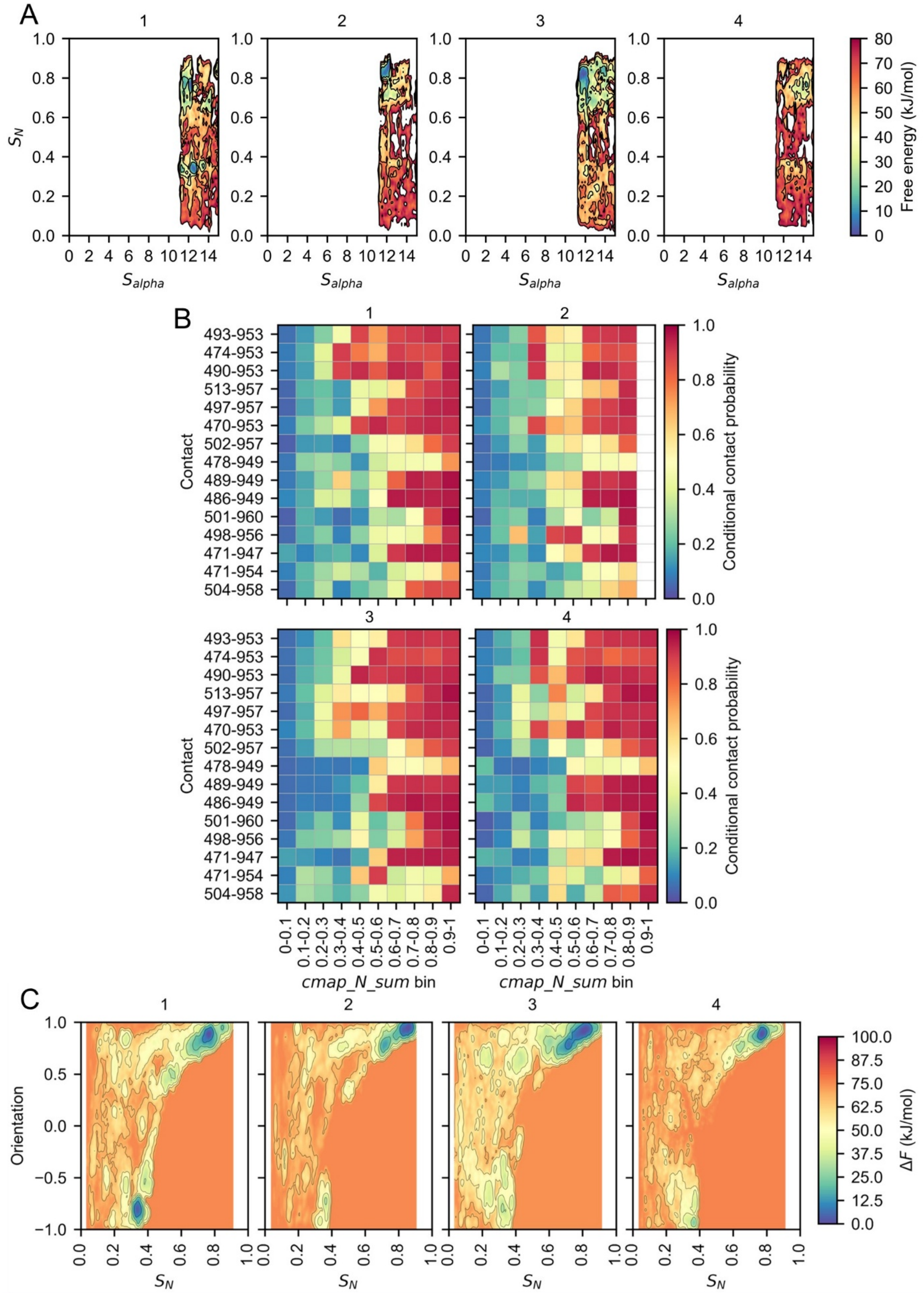


*Figure S14. Mechanism of folding and binding for the helix restricted data. (**A**) Reweighted FES projected to $S_{alpha}$ and $S_N$ coordinates for each independent OneOPES repeat. (**B**) Two-dimensional conditional contact-probability maps for each independent OneOPES repeat, calculated from the reweighted*

*ensemble. The x-axis shows ranges of the $S_N$ CV, while the y-axis lists the residue pairs forming interchain contacts. (**C**) Reweighted FES projected to $S_N$ and orientation coordinates for each independent OneOPES repeat.*
